# Regulatory architecture and evolution of the triphasic life cycle in the red alga *Bostrychia moritziana*

**DOI:** 10.64898/2026.11.10.765333

**Authors:** Romy Petroll, Prakash Sivakumar, Valentin Bernard, John A. West, Susana M. Coelho, Michael Borg

## Abstract

The triphasic life cycle of florideophyte red algae represents the most complex life cycle among photosynthetic eukaryotes, yet its developmental and evolutionary origins remain unresolved. Here, we generated a comprehensive transcriptomic atlas of *Bostrychia moritziana*, a model red alga with a highly elaborate triphasic life history. We show that cellular specification across generations, reproductive structures and unicellular reproductive cell types in *Bostrychia* is associated with discrete regulatory modules that frequently cluster within the genome. We reveal that development of the cystocarp, the reproductive structure that houses the additional diploid phase characteristic of the triphasic life cycle, involves extensive and coordinated post-fertilisation reprogramming of the zygote-derived carposporophyte and surrounding maternal tissue. Evolutionary transcriptomics support a scenario in which triphasy evolved through successive elaboration of ancestral developmental programs, with lineage-specific genes and expanded transcription factor families progressively integrated into its regulatory architecture. Together, our findings provide a molecular framework for understanding the origin and evolution of the triphasic life cycle, offering broader insights into how increasingly complex multicellular life cycles emerged across eukaryotes.

## Main

The alternation of generations, in which multicellular haploid and diploid life forms develop within a single life cycle, has evolved independently across multiple eukaryotic lineages^1,2^. Despite its profound evolutionary and ecological consequences^3–5^, the molecular basis of how these distinct generations are developmentally established and differentiated remains poorly understood outside a few well-studied systems, particularly bryophytes, angiosperms and brown algae^6–10^. The least understood haploid-diploid life cycles are those of the red algae (Rhodophyta), an ancient and highly diverse phylum of Archaeplastida that share a common ancestry with land plants^11–13^. Red algae independently evolved complex multicellularity and an alternation of generations^14^, making them a powerful comparative system to investigate the origin and diversification of complex eukaryotic life cycles.

The Florideophyceae, which comprise ∼95% of extant red algal diversity, evolved an elaborate life cycle involving haploid male and female gametophytes followed by two successive diploid generations - the zygote-derived, maternally-provisioned carposporophyte and the free-living tetrasporophyte^15,16^ (**Fig. 1**). Phylogenetic studies suggest that this so-called triphasic life cycle arose during the Neoproterozoic soon after the emergence of the Florideophyceae, approximately 943-661 million years ago, where it has remained remarkably conserved throughout radiation of the group^12^. Although some earlier diverging lineages have secondarily simplified this pattern, triphasy is otherwise near universal among the Florideophyceae and is thought to be a key innovation underpinning their extraordinary diversification^12^.

**Figure 1.**
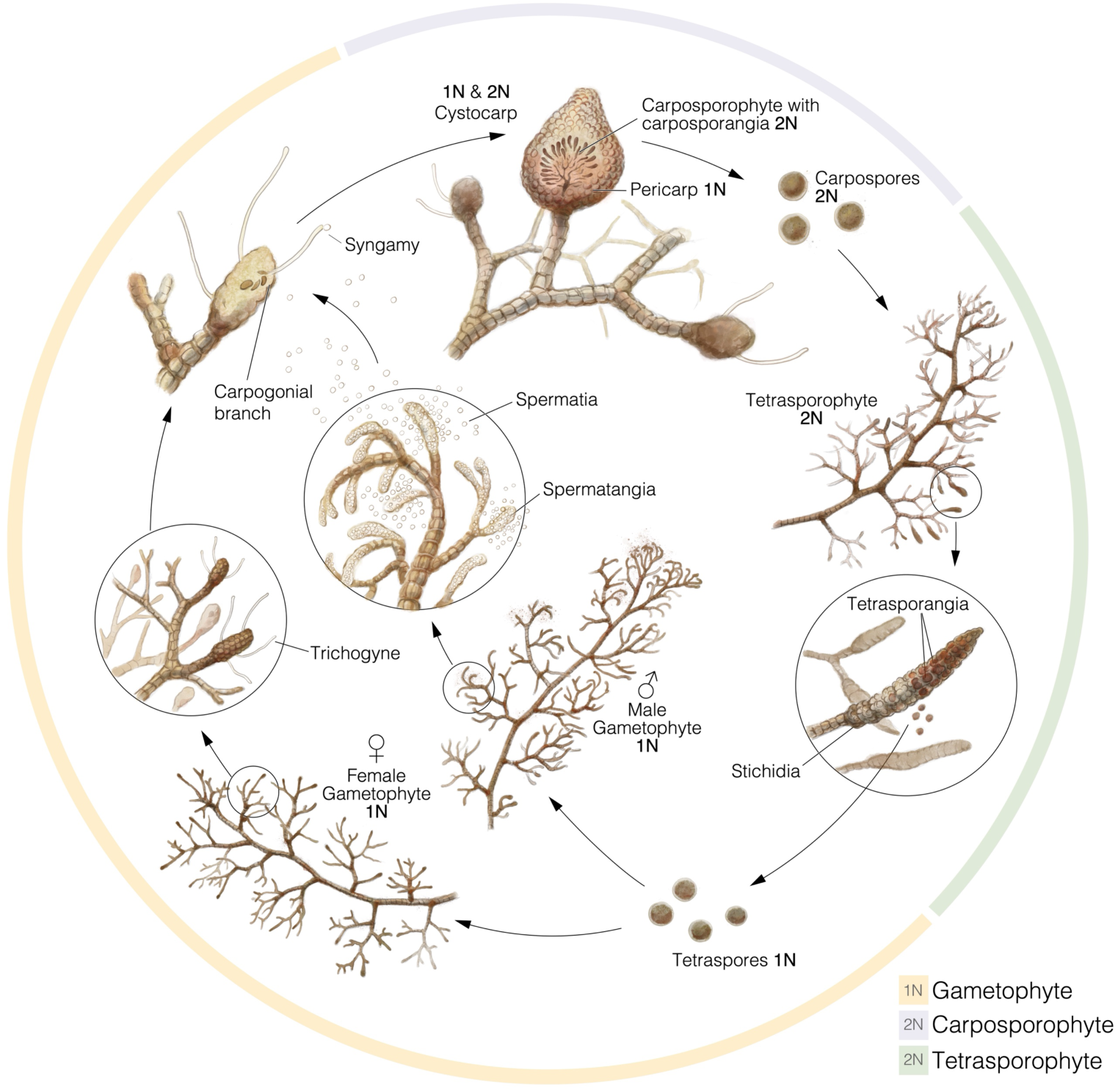
The triphasic life history of *Bostrychia moritziana*. The three successive multicellular generations in the truphasic life cycle are the haploid male and female gametophytes (yellow outline), a diploid carposporophyte retained on the female gametophyte (purple outline), and a free-living diploid tetrasporophyte (green outline). The cycle begins with the haploid gametophytes. Female gametophytes develop carpogonial branches (procarps) at their tips, bearing a receptive hair-like projection called the trichogyne. Male gametophytes produce spermatangia that release non-motile spermatia. Following attachment of a spermatium to the trichogyne and fertilisation of the carpogonium, the resulting diploid zygote develops into a multicellular carposporophyte. This remains attached to the female gametophyte and becomes enclosed by a protective pericarp derived from haploid maternal tissue, together forming the cystocarp. At maturity, the carposporophyte produces numerous diploid carpospores that are released from the cystocarp and germinate into free-living diploid tetrasporophytes. Although morphologically similar to the gametophytes, tetrasporophytes develop specialised reproductive structures called stichidia, which bear multiple tetrasporangia. Within each tetrasporangium, meiosis produces four haploid tetraspores. These are released and germinate into either male or female gametophytes, completing the triphasic life cycle.

The triphasic life cycle raises evolutionary questions beyond those posed by biphasic systems, as it requires explanation for how a second, functionally distinct diploid phase evolved and elaborated into a complex reproductive structure. Did the carposporophyte evolve as a novel post-fertilisation developmental program associated with zygote retention, maternal provisioning and zygotic amplification, through the progressive modification and partitioning of an ancestral sporophytic program, or perhaps through a combination of both processes? Resolving this question is central to understanding the evolution red algal reproduction, but also to broader questions of how complex eukaryotic life cycles arise and diversify. Despite its evolutionary significance, the regulatory basis of triphasic development remains unexplored. While existing studies have documented transcriptional reprogramming associated with different biological processes in red algae, including cell wall remodeling in *Chondrus crispus*^17^, sex-biased expression in *Bostrychia moritziana*^18,19^, and diurnal rhythms in *Gracilariopsis* ^20^, none have resolved gene expression across the triphasic life cycle in complex florideophyte red algae.

Here, we leverage genomic resources for the emerging model red alga *Bostrychia moritziana* (Sonder ex Kützing) (herein *Bostrychia*)^21^, a member of the morphologically complex Ceramiales order, to decipher regulatory and evolutionary relationships in the triphasic life cycle. Using targeted tissue dissection and low-input transcriptomics, we generate a comprehensive transcriptomic atlas spanning each major generation, reproductive structure and unicellular reproductive cell type. Using differential gene expression, co-expression networks, allele-specific expression and comparative evolutionary genomics, we show how each stage and structure across the life cycle is specified by discrete, chromosomally clustered regulatory networks enriched for distinct complements of transcription factors (TFs). We further demonstrate that the carposporophyte does not simply recapitulate the spore-producing program of the tetrasporophyte, but is rather assembled from both novel and redeployed gene networks. Finally, phylostratigraphic analyses reveal progressive recruitment of red algal lineage-specific genes into an ancestral developmental framework, providing an evolutionary scenario for the origin and elaboration of the triphasic life cycle. This study provides a comprehensive molecular view of red algal development, providing a foundation for facilitating mechanistic studies of the remarkable developmental complexity of red seaweeds.

### Transcriptional reprogramming across the *Bostrychia* life cycle

*Bostrychia* is a filamentous red seaweed that displays a fully elaborated form of the triphasic life cycle first described in the genus *Polysiphonia*^22^ (**Fig. 1**). Although the gametophyte and tetrasporophyte are macroscopically isomorphic, highly pronounced morphological differentiation occurs across its reproductive structures (**Fig. 1**). Male gametophytes produce spermatangia and non-motile spermatia, whereas female gametophytes bear multiple procarps at the tips of their branches, each containing a carpogonium (egg cell) that receives the male nucleus through a specialised projection called the trichogyne^23–25^. Fertilisation gives rise to a diploid carposporophyte that remains intimately associated with the female gametophyte and is enclosed by maternal haploid pericarp tissue within a structure called the cystocarp^15^. The maternally-provisioned cystocarp amplifies the zygote into numerous diploid carpospores^26^. Once released, these carpospores germinate into a free-living diploid tetrasporophyte, which undergoes meiosis in tetrasporangia borne on stichidia to produce haploid tetraspores, regenerating the male and female gametophyte generations.

We first improved our previously published *Bostrychia* genome assembly by generating additional long-read sequencing data to close assembly gaps and refine gene annotation (**Extended Data Fig. 1a-d**; **Supplemental Table 1**)^19^. Using tissue dissection and low-input transcriptomics, we then generated transcriptomes across 13 developmental stages: immature and mature cystocarps, carposporophyte tissue, pericarp, carpospores, tetrasporophyte thallus tissue, stichidia, tetraspores, female thallus tissue, female tips bearing procarps, male thallus tissue, spermatangia and spermatia (**Fig. 2a-g; Supplemental Table 2**). Reproductive structures were removed from the three thallus samples to distinguish vegetative from reproductive tissues, while the embedded carposporophyte tissue was released from the cystocarp to profile it separately from its surrounding pericarp layer.

**Figure 2.**
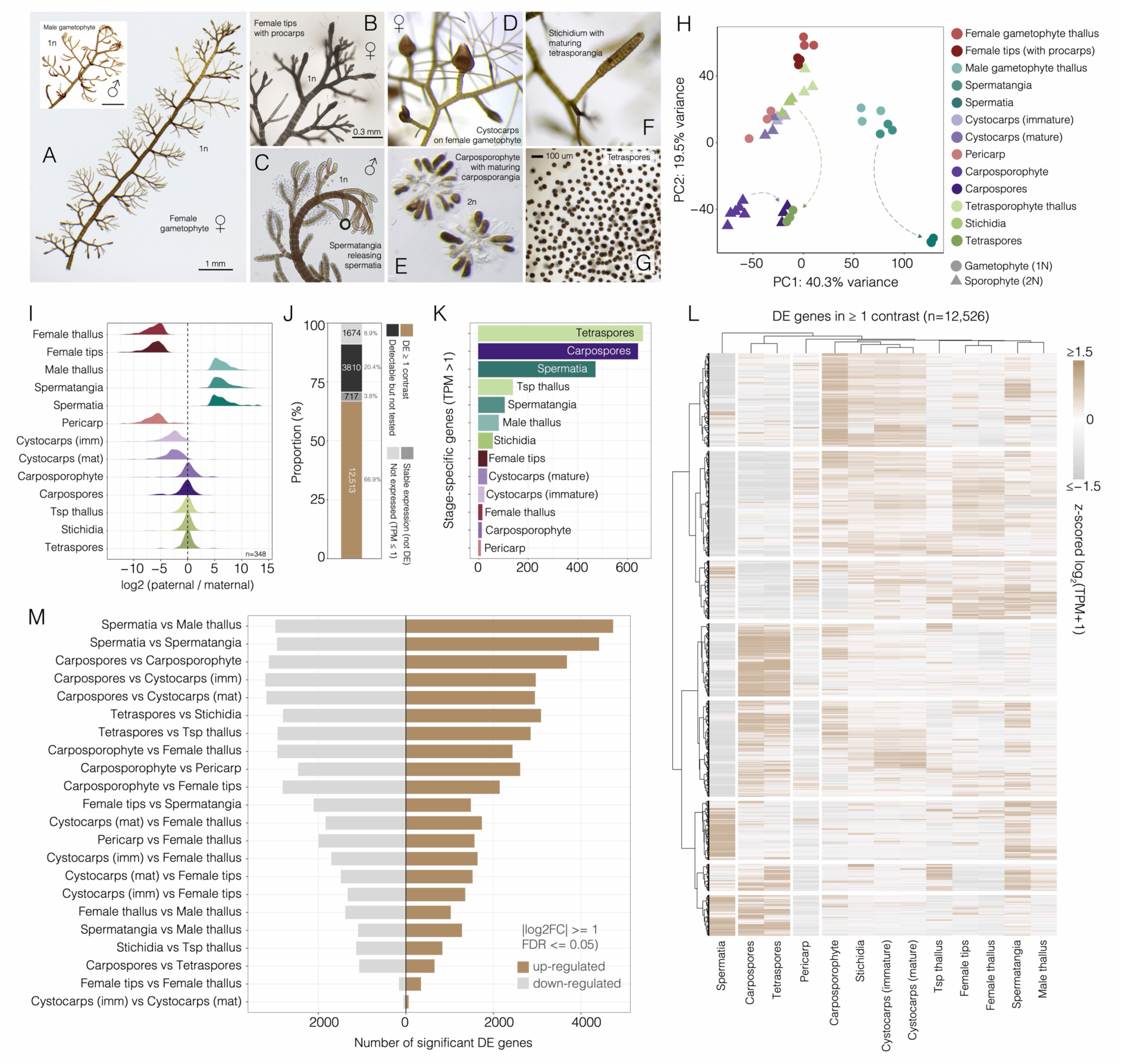
Reproductive development and transcriptional differentiation across the triphasic life cycle of *Bostrychia moritziana*. (A–G) Representative images of the reproductive generations, tissues and cells sampled for transcriptomic analysis. (A) Female and male haploid gametophytes. (B) Female gametophyte tips bearing procarps. (C) Male gametophyte branches bearing spermatangia that release spermatia. (D) Cystocarps developing on the female gametophyte. (E) Dissected diploid carposporophyte containing developing carposporangia. (F) Stichidium containing maturing tetrasporangia on the diploid tetrasporophyte. (G) Released tetraspores. (H) Principal component analysis (PCA) of transcriptomic profiles across reproductive stages and tissues. Colours indicate sample identity, while circles and triangles distinguish haploid gametophytic and diploid sporophytic samples, respectively. Dashed arrows indicate transitions between a multicellular stage and their corresponding unicellular reproductive stages. (I) Distributions of paternal-to-maternal allelic expression ratios across developmental stages and tissues. Ratios were calculated as log₂((paternal counts + 0.5)/(maternal counts + 0.5)) for genes with at least ten allele-informative counts. The total number of genes with informative SNP differences varied across stages. (J) Proportions of genes classified as differentially expressed in at least one comparison, stably expressed, not expressed and not tested. (K) Number of stage-specific genes (mean TPM > 1) in each of the assayed stages. (L) Hierarchically clustered heatmap of genes differentially expressed in at least one pairwise comparison (n = 12,526). Z-score normalised log₂(TPM + 1) expression across reproductive stages and tissues is shown. (M) Numbers of significantly upregulated (brown) and downregulated (grey) genes in selected pairwise comparisons. Differential expression was defined by an absolute log₂ fold-change ≥ 1 and a false discovery rate (FDR) ≤ 0.05. Imm, immature; mat, mature; Tsp, tetrasporophyte; DE, differentially expressed; TPM, transcripts per million.

Principle component analysis (PCA) and sample–sample correlation analyses confirmed high reproducibility among replicates, which clearly clustered by biological identity, while gene detection and sensitivity remained high despite low RNA input (**Fig. 2h; Extended Data Fig. 2a-d**). Female and tetrasporophyte samples were more similar to one another than to male samples, suggesting partially shared developmental programs (**Fig. 2h**). Male samples occupied a distinct transcriptomic space, with reproductive transitions introducing further separation, particularly among unicellular stages (**Fig. 2h**). Carposporophyte samples were clearly distinct from both cystocarp and pericarp samples, consistent with the latter representing composite, maternally-dominated transcriptomes (**Fig. 2h**). Allele-specific expression supported this interpretation. Using informative SNPs that distinguished maternal and paternal alleles, the pericarp showed strong maternal bias similar to the female gametophyte samples, whereas as the carposporophyte showed near balanced allelic expression characteristic of the tetrasporophyte stages (**Fig. 2i**). Carpospores and tetraspores clustered closely but remained distinct, whereas spermatia were sharply separated from spermatangia (**Fig. 2h**). Together, these data support the purity and reproducibility of our transcriptomic datasets and show that the largest transcriptomic changes accompany differentiation into unicellular stages, whereas multicellular stages exhibit more gradual transcriptional change.

Global expression analyses further revealed extensive transcriptional reprogramming across the life cycle (**Fig. 2j,l; Supplemental Table 3**). Most protein-coding genes were detected and showed stage-dependent changes in expression (TPM > 1; 12,513 of 18,714; 66.9%), whereas only a small fraction of genes were stably expressed (717; 3.8%) or not detected (TPM ≤ 1; 1,674; 8.9%) (**Fig. 2j**). A further 3,801 expressed genes fell below our count threshold for differential expression analysis (see Methods). The largest number of differentially-expressed genes (DEGs) were observed in transitions to spermatia and spores (several thousands), whereas maturation steps, such as immature to mature cystocarps, involved relatively modest changes (**Fig. 2m; Extended Data Fig. 3**). Despite this, most genes (14,732; 78.7%) were expressed across multiple stages, suggesting broadly shared transcription throughout the life cycle (**Extended Data Fig. 4a**). These broadly expressed genes spanned a broad dynamic range, suggesting that developmental transitions involve quantitative modulation of gene expression rather than strict on-off regulation (**Extended Data Fig. 4b**). Genes were generally expressed at higher levels in spermatangia and carpospores relative to their average across stages, whereas spermatia showed lower expression and had the fewest detectable genes (**Extended Data Fig. 4c,d**). Stage-specific genes accounted for only a small fraction of the transcriptome (2,308; 12.3%), with the majority concentrated in spores and spermatia (**Fig. 2k; Extended Data Fig. 4a; Supplemental Table 3**). Both the proportion of stage-dependent genes and the genome-wide range of expression were similar when normalisation size-factors were estimated using the 724 least variable genes **(Extended Data Fig. 4e)**. Stage-dependent expression also persisted when testing was restricted to this low-variance subset (**Extended Data Fig. 4f**), further supporting the robustness of the developmental expression signals. Together, these results show that developmental transitions in *Bostrychia* involve dynamic modulation of a shared transcriptional program, with stage identity associated with relatively small, specialised sets of genes.

### Regulatory architecture of generation and sex identity in *Bostrychia*

To investigate the transcriptional basis of generation identity across the triphasic life cycle, we focused on genes consistently upregulated (*generation-bia*sed) or downregulated (*generation-repressed*) across the three multicellular generations. Spermatia and spores were excluded to distinguish generation-associated expression from terminal differentiation programs.

Overall, 12.3% of protein-coding genes (2,295 of 18,714 genes) showed generation-biased expression, with 916 genes gametophyte-biased, 869 carposporophyte-biased, and 510 tetrasporophyte-biased (**Fig. 3a,b; Extended Data Fig. 5a; Supplemental Table 3**). Among these, the proportion expressed exclusively in the corresponding generation was highest in gametophytes (22%; 200/916), followed by the tetrasporophyte (15%; 79/510) and carposporophyte (10%; 84/869) (**Fig. 3b**). Conversely, 13.8% of protein-coding genes (2,581 of 18,714) showed generation-specific repression, most prominently in the carposporophyte (1,863 genes), compared with the gametophyte (460) and tetrasporophyte (258) (**Fig. 3a,b; Extended Data Fig. 5b; Supplemental Table 3**). Among these repressed genes, expression fell below the detection threshold (TPM < 1) in 38% of carposporophyte-repressed and 26% of tetrasporophyte-repressed genes, but only 6% of gametophyte-repressed genes (**Fig. 3b**). These observations further support a model in which major life cycle transitions involve selective activation and repression of a relatively small subset of genes against a backdrop of broadly shared gene expression.

**Figure 3.**
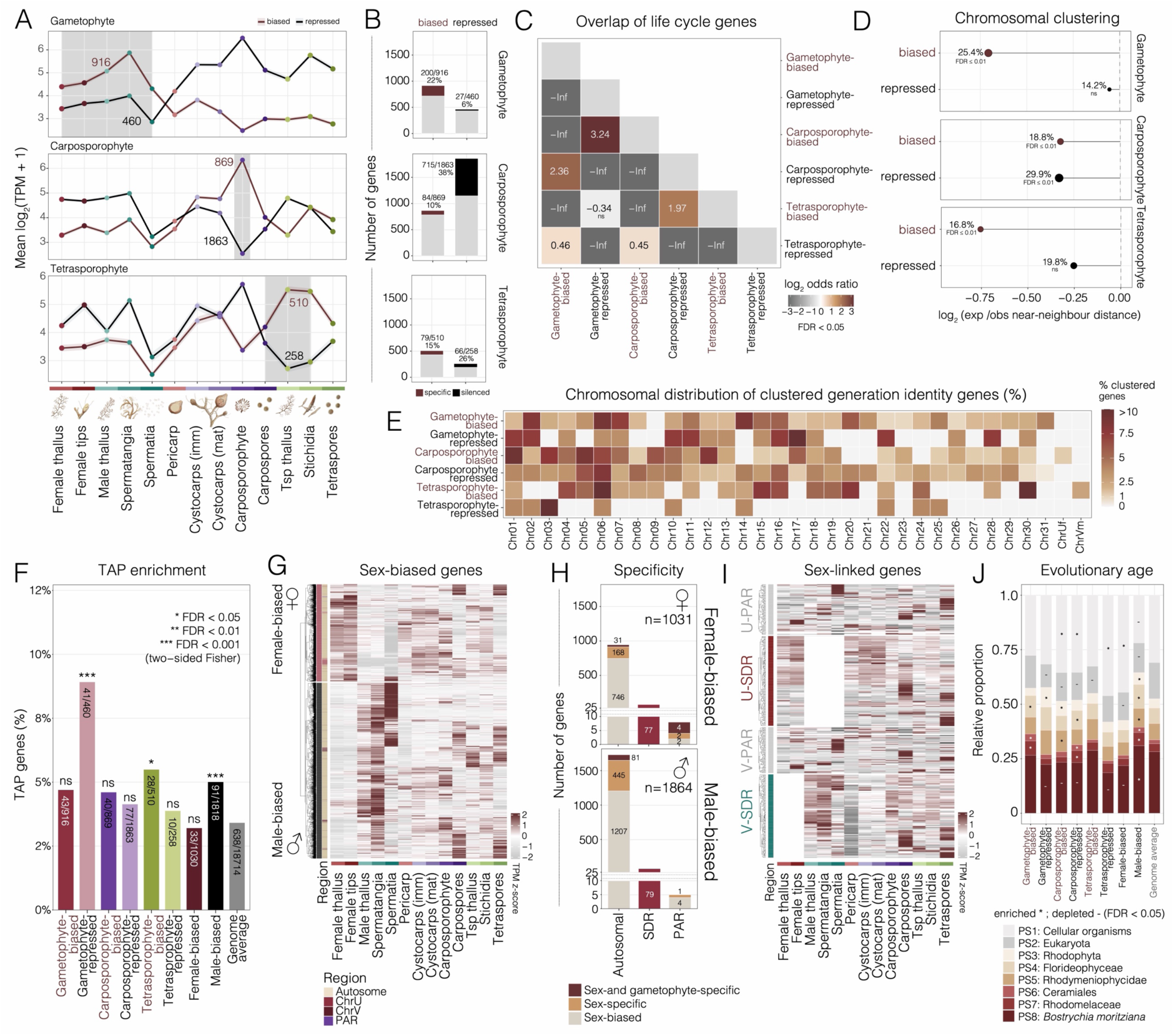
Generation- and sex-specific transcription across the *Bostrychia* triphasic life cycle. (A) Mean log₂(TPM + 1) expression profiles of generation-biased and generation-repressed genes across reproductive stages and tissues. The numbers of genes in each set is indicated, with the selected stages defining the corresponsding gene set highlighteds with grey shading. (B) Number and proportion of generation-biased and generation-repressed genes in gametophytes, carposporophytes and tetrasporophytes. (C) Pairwise overlap between generation-biased and generation-repressed gene sets, expressed as log₂ odds ratios. Significant associations (FDR < 0.05) are highlighted, while non-significant associations are shown in grey. (D) Chromosomal clustering of generation-biased and generation-repressed genes was assessed by comparing observed distances between neighbouring genes with expected distances. oints indicate log₂(observed/expected median nearest-neighbour distance), with negative values indicating shorter distances than expected from chromosome-stratified permutations. The proportion of genes in proximity blocks and FDR-adjusted significance levels are indicated. (E) Chromosomal distribution of clustered generation-biased and generation-repressed genes, expressed as the percentage of genes within each chromosome belonging to the corresponding clustered gene set. (F) Proportion of transcription-associated protein (TAP) genes among generation- and sex-biased or repressed gene sets, compared with the genome-wide average. Enrichment significance was assessed using two-sided Fisher’s exact tests with FDR correction. (G) Hierarchically clustered heatmap of female- and male-biased genes across reproductive stages and tissues. The annotation strip indicates autosomal, sex-linked and pseudoautosomal locations. (H) Numbers of female- and male-biased genes located on autosomes, sex-determining regions (SDRs) and pseudoautosomal regions (PARs), classified as sex-biased, sex-specific or both sex- and gametophyte-specific. (I) Heatmap showing expression profiles of genes located in the U- and V-linked PARs and SDRs across reproductive stages and tissues. In (G) and (I), colours indicate row-scaled TPM expression values (z-scores). (J) Relative proportions of genes assigned to successive evolutionary phylostrata within generation- and sex-associated gene sets. Asterisks indicate significant enrichment and dots indicate significant depletion (FDR < 0.05). Phylostrata range from broadly conserved cellular genes (PS1) to genes specific to *B. moritziana* (PS8). Imm, immature; mat, mature; Tsp, tetrasporophyte; TPM, transcripts per million; FDR, false discovery rate.

Overlap patterns among generation-biased and generation-repressed genes (i.e. generation identity genes) revealed distinct regulatory shifts across the triphasic life cycle. Gametophyte-biased genes significantly overlapped with genes repressed in both diploid generations, suggesting selective repression of the gametophytic program during sporophytic development (**Fig. 3c**). Conversely, carposporophyte-biased genes were enriched among those repressed in both the gametophyte and tetrasporophyte, whereas tetrasporophyte-biased genes overlapped primarily with carposporophyte-repressed genes, but not those repressed in the gametophyte (**Fig. 3c**). These patterns suggest that, rather than sharing a common sporophyte-specific expression program, the two diploid generations exhibit distinct transcriptional identities, with carposporophyte development involving selective repression of tetrasporophyte-biased genes. Allele-specific expression further supported post-fertilisation reprogramming within the cystocarp, revealing nearly equal maternal and paternal transcriptional contributions in the carposporophyte (**Fig. 2i**). Although the surrounding pericarp remained strongly maternally biased (**Fig. 2i**), its transcriptional profile more closely resembled that of the cystocarp than the female gametophyte (**Fig. 2h**), indicating substantial transcriptional reprogramming of maternal gametophytic tissue following fertilisation. Together, these findings suggest that cystocarp development involves distinct but coordinated transcriptional reprogramming of the carposporophyte and surrounding maternal tissues, rather than the simple re-deployment of a shared sporophytic program within the female gametophyte.

Four of the six generation identity gene sets also exhibited non-random genomic organization, clustering significantly more than expected by chance (**Fig. 3d-e; Extended Data Fig. 5c; Supplemental Table 4).** The strongest clustering was observed among gametophyte-biased and tetrasporophyte-biased genes, whereas carposporophyte-repressed genes formed the greatest number of clusters and had the highest proportion of genes within proximity blocks, reflecting the size and local organisation of this gene set (**Fig. 3d**). Generation identity genes were also associated by distinct repertoires of transcription-associated proteins (TAPs), with gametophyte-repressed and tetrasporophyte-biased genes showing particularly strong enrichment (**Fig. 3f; Extended Data Fig. 5a-b**). Triphasic generation identity is thus shaped by locally organised genomic architecture that like facilitates coordinated gene regulation by phase-specific regulatory proteins. Notably, similar TAP families, including C2H2 zinc-finger and MADS-box TFs, were represented across generation-specific gene sets, suggesting that regulatory specialisation may have involved functional diversification of TFs within shared families. Together, these findings suggest that generation identity is associated with both local genomic clustering and the differential deployment of regulatory proteins across the triphasic life cycle.

Because gametophyte-biased genes were defined using both male and female samples, we next examined sex-biased gene (SBG) expression across the two gametophytes (**Fig. 3g; Supplemental Table 3**). We identified 1,031 female-biased genes (FBGs) and 1,864 male-biased genes (MBGs), of which 1,030 FBGs and 1,818 MBGs were reliably detected (TPM ≥ 1) in at least one sex. Within these subsets, 282 (27.4%) genes were female-specific and 606 (33.3%) male-specific (**Fig. 3h**). Only 35 FBGs (3.4%) and 82 MBGs (4.5%) were exclusively expressed in their respective gametophytes and undetectable in all other life cycle stages, indicating that most SBGs are also expressed elsewhere in the life cycle (**Fig. 3g,h**). Similarly, genes within the sex-determining regions (SDR) of the U and V were expressed not only in their respective gametophyte but also in the carposporophyte, tetrasporophyte and their respective spores, suggesting potential functions beyond gametophytic sexual differentiation (**Fig. 3i; Extended Data Fig. 5d**). The recombining pseudoautosomal region (PAR) contributed relatively few SBGs, whereas autosomes accounted for a substantial proportion, including 199 female-specific and 526 male-specific genes (**Fig. 3h**). Notably, only MBGs were significantly enriched for TAPs relative to the genome average (**Fig. 3f**), pointing to prominent regulatory reconfiguration during male gametophyte development. Together, these findings indicate that sexual differentiation in *Bostrychia* is associated with both UV sex chromosome inheritance and extensive sex-biased expression of autosomal genes.

Finally, we examined the evolutionary age and functional composition of the generation and sex identity genes, revealing distinct, non-random phylostratigraphic distributions (**Fig. 3j; Supplemental Table 3**). Red algal-specific genes were particularly enriched among gametophyte- and carposporophyte-biased and -repressed genes, suggesting contribution of lineage-specific innovations to their developmental specialisation. In contrast, *Bostrychia*-specific genes were significantly depleted across most identity genes, with MBGs representing a notable exception given their enrichment for younger phylostrata (**Fig. 3j**), mirroring the rapid evolution of male reproductive genes observed across eukaryotes^27–29^. Functional enrichment revealed differences among generation identity and sex-biased genes (**Supplemental Table 5**), including functions related to heat shock proteins, cell cycle regulation, chromosome and chromatin organization, and extracellular matrix and cell surface components. However, at least half of these genes are specific to red algal lineages and remain poorly annotated, likely biasing these results towards deeply conserved genes with well-characterised functions. Together, these findings reveal distinct evolutionary and functional signatures of generation- and sex-associated transcriptional specialisation, which draws on both evolutionary conserved genes and red algal lineage-specific innovations.

### Discrete co-expression networks underpin reproductive development in *Bostrychia*

To further resolve the regulatory programs associated with cell fate specification in *Bostrychia*, we identified genes that distinguish individual developmental stages, reproductive structures and cell types. Marker gene analysis recovered 609 with highly enriched diagnostic expression, predominantly in male reproductive stages like spermatia (236), but also in carpospores (106) and tetraspores (69) (**Fig. 4a**). Significant functional enrichment was largely restricted to spermatial markers, which were associated with cell wall and polysaccharide metabolism, protein kinase signalling and cyclin-related functions (**Supplemental Table 5**). Several TFs also showed stage-specific enrichment, including an RKD factor in procarp-bearing female tips, raising intriguing parallels with the role of this TF family in female reproductive cell specification across land plants^30–32^.

**Figure 4.**
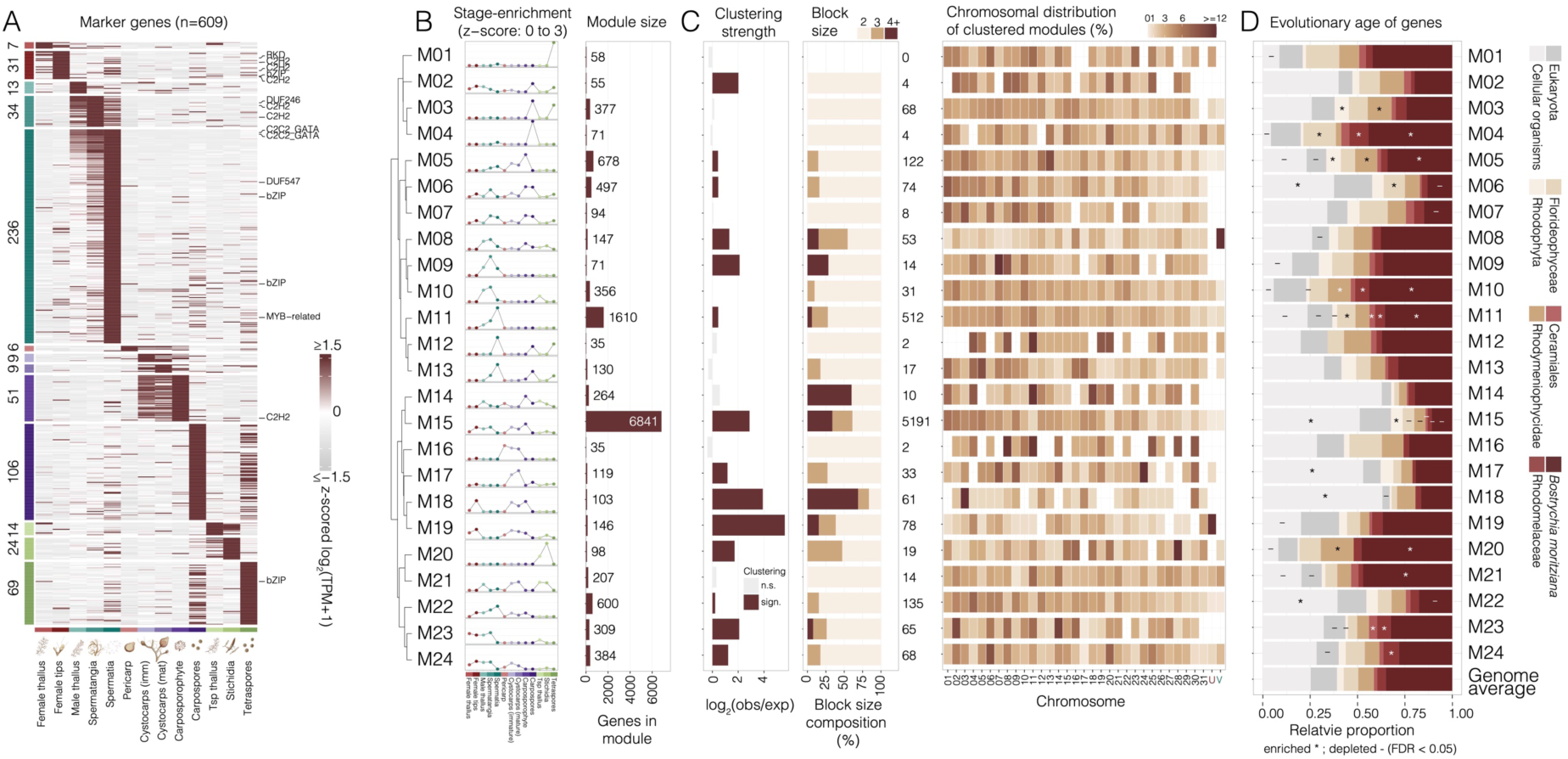
Spatial organisation and evolutionary features of co-expression modules in *Bostrychia*. **(A)** Heatmap showing the expression profiles of 609 stage-associated marker genes across reproductive stages and tissues. Genes are grouped according to their stage-associated expression patterns, with group sizes indicated on the left and selected TF families annotated on the right. Colours represent z-score normalised log₂(TPM + 1) expression values. **(B)** Hierarchical clustering of 23 gene co-expression modules and one unassigned class (M21), showing their relative expression enrichment across developmental stages and tissues (z-scores, 0–3) and the number of genes assigned to each module. Module M21 represents an unassigned class and thus does not represent a coherent co-expression module. **(C)** Genomic organisation of co-expression modules. Clustering strength is expressed as log₂(expected/observed median nearest-neighbour distance), with positive values indicating shorter observed distances than expected. The expected distance is the median of the permutation-derived median distances. Block-size composition indicates the proportion of clustered genes occurring in blocks of two, three or at least four genes. The heatmap shows the chromosomal distribution of clustered genes for each module, expressed as the percentage of clustered module genes on each chromosome. **(D)** Evolutionary age composition of genes within each co-expression module based on phylostratigraphic assignments. Stacked bars indicate the relative proportion of genes assigned to successive evolutionary strata, from broadly conserved cellular genes (PS1) to *Bostrychia*-specific genes (PS8), alongside the genome-wide average. Asterisks and dashes indicate significant enrichment and depletion, respectively (FDR < 0.05). Imm, immature; mat, mature; Tsp, tetrasporophyte; TPM, transcripts per million; FDR, false discovery rate.

Weighted gene co-expression analysis provided a broader view of developmental regulation, assigning 13,078 genes to 23 modules that collectively distinguished all major stages of the life cycle, with only 207 genes remaining unassigned in module M21 (**Fig. 4b; Supplemental Table 3**). Larger modules were broadly expressed across multicellular gametophytic and sporophytic tissues, whereas smaller modules were tightly associated with individual reproductive structures, including spermatangia, spermatia, the carposporophyte, stichidia and spores (**Fig. 4b**). TFs were distributed non-randomly across modules, with candidate regulatory hubs concentrated in reproductive tissue- and cell type-enriched modules (**Extended Data Fig. 6**), suggesting that developmental specialisation is coordinated by distinct combinations of transcriptional regulators.

Given that the carposporophyte ultimately defines the triphasic life cycle, we examined its associated modules more closely. M05 was the only module strongly enriched in the carposporophyte, whereas the remaining associated modules (M06, M07, M14 and M15) were shared with other developmental stages. However, comparisons of network connectivity across these five modules revealed a distinct carposporophyte-associated organization, rather than one resembling the tetrasporophyte or its reproductive stichidia (**Extended Data Fig. 7**). Although the carposporophyte and stichidia share spore-producing functions and several co-expression modules, including M14 and M15, their connectivity patterns differed substantially, with stichidia showing weaker correlation to the carposporophyte than several other developmental stages. These findings suggest that the carposporophyte represents a distinct transcriptional state characterised by the differential organisation of developmental modules shared across the life cycle, including those active in the tetrasporophyte.

Given the genomic clustering of generation identity genes (**Fig. 3**), we next asked whether co-expression modules had similar spatial organisation. Chromosomal clustering was widespread, with 14 of 24 modules significantly more clustered within the genome than expected by chance (**Fig. 4c; Supplemental Table 4**). Several formed pronounced chromosomal hotspots, most notably the female tip–cystocarp module M18, which concentrated within a large tandemly repeated HSP20 gene cluster on chromosome 3 (**Fig. 4c; Supplemental Table 4**), a gene family previously implicated in female gametophyte development in red algae^33^. Modules associated with procarp-bearing tips and the carposporophyte (M19), and with spermatangia (M08), were enriched on the U and V sex chromosomes, respectively (**Fig. 4c**). Their strongest TF association included the corresponding U- and V-linked TALE-HD TFs (**Extended Data Fig. 6**). In contrast, larger and more broadly expressed modules, including M11 and M15, were distributed across multiple multiple genomic regions rather than concentrated in discrete chromosomal hotspots (**Fig. 4c**).

Phylostratigraphic analysis revealed distinct evolutionary signatures across co-expression modules, reflecting the integration of ancient and progressively younger genes into reproductive development (**Fig. 4d**). Modules enriched for deeply conserved genes (PS1– PS2) generally showed the strongest functional annotation, including the broadly expressed module M15, associated with core cellular functions, and modules active in procarp-bearing female tips and the carposporophyte (M06, M17 and M18), enriched for DNA metabolism, chromosome organisation, cell division, carbonic anhydrase and heat shock protein functions (**Fig. 4d; Supplemental Table 5**). Other ancient-enriched modules included M22, which showed strong spermatial expression and enrichment for RNA polymerase-associated functions. Modules enriched for genes originating during Rhodophyta, Florideophyceae and Rhodymeniophycidae evolution were more closely associated with reproductive specialisation. These included the spore-specific module M03, linked to extracellular proteoglycan metabolism and morphogenesis, and carposporophyte- and carpospore-associated modules M05–M07, enriched for DNA replication and repair, cell cycle progression and histone kinase-related functions (**Fig. 4d; Supplemental Table 5**).

More recently evolved genes, originating within Ceramiales and Rhodomelaceae, were particularly enriched in reproductive modules, including M11, associated with polysaccharide metabolism and CDK-like activity in male reproductive stages, and M23 and M24, associated with spermatangia and procarp-bearing female tips and enriched for extracellular matrix and VWA–collagen-like domains (**Fig. 4d; Supplemental Table 5**). The youngest, *Bostrychia*-specific genes were likewise concentrated in reproductive modules associated with the carposporophyte and carpospores (M04, M05), male reproductive stages (M10, M11) and stichidia (M20). Most of these younger modules lacked informative functional annotation, limiting interpretation of their biological roles.

Together, these findings reveal an evolutionarily layered transcriptional architecture underlying reproductive development in *Bostrychia*, in which discrete, frequently clustered co-expression modules combine deeply conserved cellular functions with lineage-specific genes. The carposporophyte exemplifies how distinct developmental states may arise through the differential deployment and reorganisation of regulatory modules shared across the triphasic life cycle.

### Evolution of developmental programs across the triphasic life cycle

To investigate how genes of different evolutionary ages contribute to the triphasic life cycle, we applied transcriptome age index (TAI) analysis, which estimates the average evolutionary age of expressed genes weighted by their transcript abundance^34^. TAI varied significantly across development (*p* = 3 × 10^−14^), revealing a non-random distribution of transcriptomic ages (**Fig. 5a**). The carposporophyte and cystocarp had the “oldest” transcriptomes (lowest TAI), whereas male reproductive stages were the “youngest” (highest TAI), particularly in spermatia.

**Figure 5.**
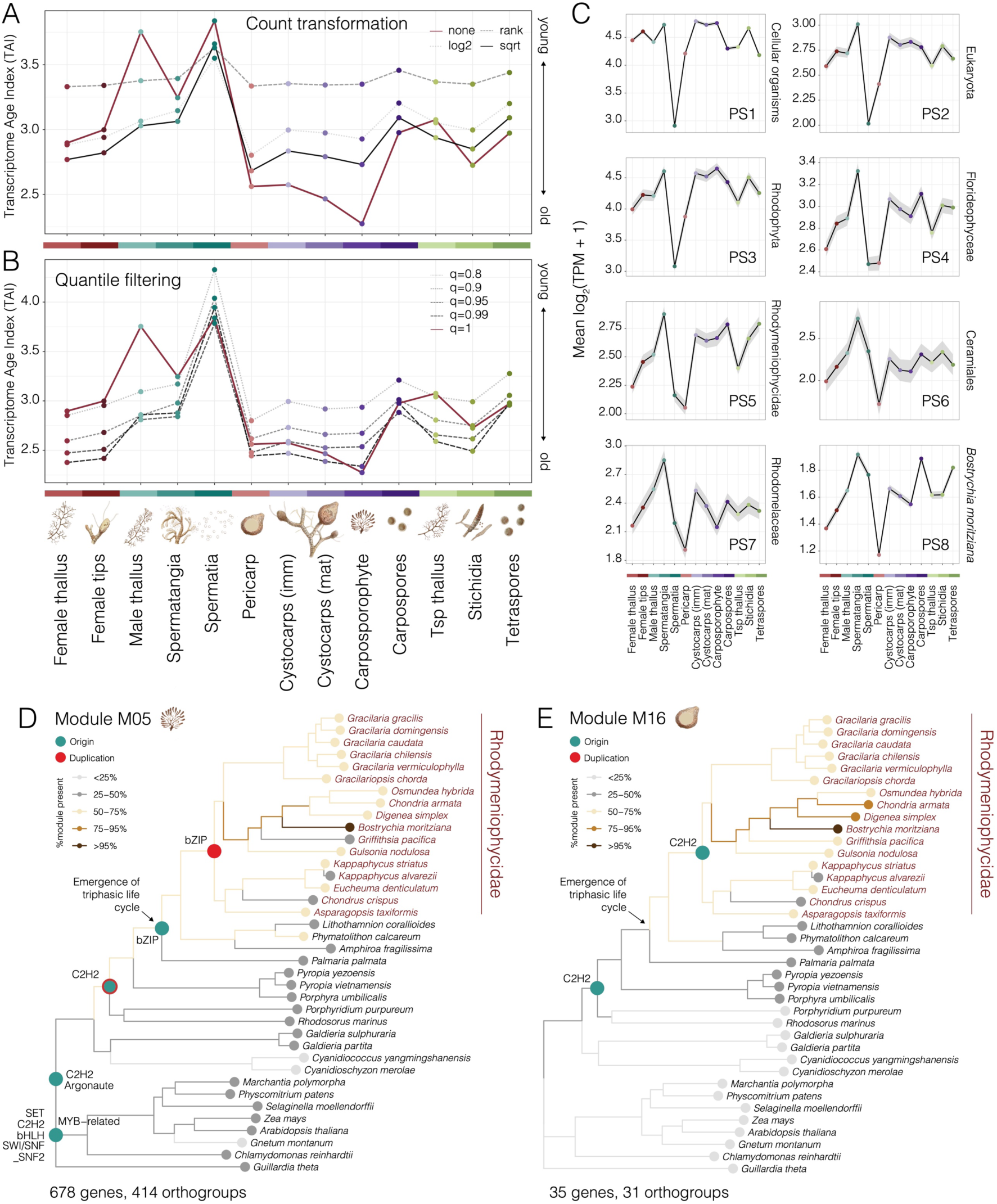
Dynamics of transcriptome evolution across the *Bostrychia* triphasic life cycle. (A) Transcriptome age index (TAI) across reproductive stages and tissues, calculated using different transformations of gene expression values (untransformed, log₂, rank and square-root). Higher TAI values indicate a greater contribution of evolutionarily younger genes to the transcriptome. (B) Sensitivity of TAI profiles to removal of genes above selected quantiles of mean expression (q = 0.8, 0.9, 0.95, 0.99 and 1). (C) Mean log₂(TPM + 1) expression profiles of genes assigned to eight evolutionary phylostrata (PS1–PS8), ranging from broadly conserved cellular genes (PS1) to *Bostrychia*-specific genes (PS8). Shaded regions indicate variation around the mean. (D, E) Phylogenetic distribution and inferred evolutionary histories of orthogroups associated with the carposporophyte-enriched module M05 (D; 678 genes, 414 orthogroups) and pericarp-enriched module M16 (E; 35 genes, 31 orthogroups). Branch colours indicate the proportion of module-associated orthogroups present in each lineage, while coloured nodes indicate inferred origins (teal) and duplications (red) of selected TF families. The inferred emergence of the triphasic life cycle is indicated on the phylogeny^12^. Imm, immature; mat, mature; Tsp, tetrasporophyte; TPM, transcripts per million.

To assess the sensitivity of these patterns to highly expressed genes, we repeated the analysis using square-root, log₂ and rank-based transformations of transcript abundance. Across transformations, female tissues, particularly the pericarp, shifted towards older transcriptomic ages, whereas the carposporophyte shifted modestly towards a younger age while remaining among the oldest transcriptomes (**Fig. 5a**). Expression quantile filtering produced similar results, suggesting that highly expressed young genes disproportionately influence TAI in female tissues, whereas the exceptionally low untransformed TAI of the carposporophyte reflects the contribution of highly expressed ancient genes (**Fig. 5b**). Despite these quantitative shifts, the overall developmental pattern was largely preserved, with female reproductive tissues and the carposporophyte generally exhibiting older transcriptomes than male reproductive stages, and the tetrasporophyte occupying an intermediate position. Unicellular reproductive cells, particularly spermatia, consistently showed relatively young transcriptomic profiles.

Stratification of transcript abundance by phylostratum further resolved evolutionary contributions to these developmental differences (**Fig. 5**). Genes originating within Rhodophyta and Florideophyceae (PS3–PS4) showed elevated expression in reproductive tissues relative to vegetative stages, particularly in spermatangia and the carposporophyte (**Fig. 5c**). During the transition from unfertilised female tips to cystocarp development, Florideophyceae- and Rhodymeniophycidae-specific genes (PS4-PS5) increased markedly, coinciding with the evolutionary elaboration of post-fertilisation development in these lineages^12^. In contrast, younger Ceramiales- and Rhodomelaceae-specific genes (PS6–PS7) were most strongly expressed in male reproductive tissues, although they also contributed to cystocarps, the carposporophyte, stichidia, spores and spermatia. The youngest, *Bostrychia*-specific genes (PS8) were predominantly expressed in spermatia and spores, account for much of their comparatively young transcriptomic profiles (**Fig. 5c**). These patterns suggest that reproductive development in *Bostrychia* draws on an ancient genetic architecture that has been elaborated by the progressive recruitment of genes from successive stages of red algal evolution. Younger genes contribute particularly strongly to male differentiation, with spores and cystocarp development also showing signatures of ongoing lineage-specific specialisation.

To investigate the evolutionary history of developmental co-expression modules, we reconstructed the ancestral presence and absence of their constituent orthogroups across the red algal phylogeny (**Extended Data Fig. 8**). This analysis revealed particularly striking patterns within co-expression modules associated with post-fertilisation development. Orthogroups represented in modules associated with the carposporophyte (M05), pericarp (M16) and carpospores (M02 and M03) were inferred to have originated early in red algal evolution and were broadly retained across Rhodymeniophycidae, including Ceramiales, where post-fertilisation development became particularly elaborate^12^ (**Fig. 5d-e**; **Extended Data Fig. 8)**. In contrast, these orthogroups showed reduced representation in Palmariales, which secondarily reverted to a biphasic life cycle, and Corallinophycidae, which lost post-fertilisation cell-to-cell fusion^12^. We further traced the evolutionary histories of the TFs associated with these modules, mapping their inferred origins and duplication events onto the ASR phylogeny (**Fig. 5d-e; Supplemental Table 6; Supplemental Table 7**). The carposporophyte-associated module M05 showed repeated expansion of key TF families. A C2H2 orthogroup was inferred to have originated near the base of multicellular red algae and subsequently duplicated within the Rhodymeniophycidae, whereas a bZIP orthogroup was inferred to have emerged around the origin of triphasy and undergone further duplication within the same lineage (**Fig. 5d**). A C2H2 orthogroup associated with the pericarp module M16 showed a similar pattern of early origin followed by lineage-specific duplication (**Fig. 5e**). Together, these findings suggest that the progressive expansion and diversification of TF families are likely to have contributed to the elaboration of post-fertilisation developmental programs during florideophyte evolution, potentially facilitating the specialisation of regulatory functions associated with cystocarp development.

## Discussion

Alternation of generations has evolved repeatedly across eukaryotes, with the triphasic life cycle of florideophyte red algae remaining among the least understood at the molecular level. Here, we investigated developmental gene expression in *Bostrychia*, a red macroalga with a highly elaborate triphasic life history. Despite pronounced morphological differences among generations, most genes were expressed across multiple developmental stages, with generation identity instead associated with relatively small sets of differentially expressed genes enriched for distinct complements of TFs. These findings suggest that developmental transitions in *Bostrychia* involve selective rewiring of regulatory networks within a broadly shared program of gene expression.

Co-expression analysis further revealed a spatially structured regulatory architecture in *Bostrychia*. Many co-expression modules, particularly those associated with reproductive development, showed significant genomic clustering. Such organisation may facilitate coordinated expression through shared *cis*-regulatory elements or local chromatin environments, although tandem duplication and the retention of neighbouring genes could also contribute to these patterns^35^. Particularly notable was the enrichment of female- and male-associated co-expression modules on the U and V sex chromosomes, respectively, which were strongly associated with sex-linked TALE-HD TFs. Although sex-liked TALE-HD factors appear to be a unique feature of red algae^19,36^, related TALE-HD regulators control life cycle transitions in green algae, bryophytes and brown algae^6–8,37–40^. More broadly, sex-linked TFs also contribute to sexual differentiation in other organisms with UV sex determination. In the liverwort *Marchantia polymorpha*, the U-linked TF BPCU acts upstream of autosomal regulators of female development^41^, whereas in the brown alga *Ectocarpus sp*, the V-linked HMG-domain factor *MIN* is implicated in male sex determination alongside extensive autosomal sex-biased expression^42,43^. These parallels suggest that sex chromosome-linked regulators repeatedly contribute to reproductive differentiation through their association with broader autosomal transcription. The numerous generation-, sex-, stage- and cell type- associated regulatory genes we have identified here provide a rich resource for dissecting the molecular basis of red algal development.

The defining feature of the triphasic life cycle is the additional diploid carposporophyte generation. Classical morphological studies established that the zygote-derived carposporophyte develops in intimate association with, and receives nutritional support from, the female gametophyte^44^. This relationship has long been considered central to the evolution of triphasy, with maternal provisioning and zygotic amplification proposed as key innovations underlying the emergence of the carposporophyte^45,46^. However, the evolutionary relationship between the carposporophyte, the free-living tetrasporophyte and the maternal gametophyte has since remained unresolved. Our allele-specific and transcriptomic analyses reveal that the carposporophyte and pericarp are not merely anatomically associated, but undergo extensive, coordinated but distinct transcriptional reprogramming following fertilisation. At the mature developmental stage we examined, the carposporophyte had near balanced maternal and paternal allelic expression, consistent with contributions from both parental genomes, whereas the pericarp remained genetically maternal despite extensive transcriptional change. This differs from the predominantly maternal control of early sporophyte development in *Marchantia*^47^, although the parental contributions during early carposporophyte development remain unknown. The cystocarp therefore comprises genetically distinct tissues undergoing coordinated developmental reprogramming, highlighting the importance of both zygotic development and maternal tissue specialisation for its establishment.

Our comparative evolutionary analyses suggest that the carposporophyte represents a distinct developmental state arising through the differential deployment of ancestral genes shared across the life cycle. The preferential expression of evolutionarily ancient genes during carposporophyte development presents an apparent paradox, given that the carposporophyte itself represents an evolutionary innovation of the florideophyte life cycle. This apparent contradiction may be reconciled by the ancient origin of the carposporophyte, potentially dating to the Neoproterozoic^12^, together with the redeployment of deeply conserved genes during its developmental elaboration. The preferential deployment of evolutionarily ancient genes echoes findings in brown algae, where ancient genes contribute more prominently to multicellular development^48^. Our observations are thus consistent with the broader evolutionary principle that morphological innovation often arises through the co-option and reorganisation of pre-existing developmental regulatory systems^49–52^. The carposporophyte and pericarp programs both contained C2H2 and bZIP TFs, families that underwent repeated expansion during red algal evolution, including within Florideophyceae and Rhodymeniophycidae^19,53^. The association of these diversified TF families, alongside the presence of lineage-specific genes that arose within these clades, suggests that both may have contributed to the increasing complexity of cystocarp development. Notably, many of the genes associated with these programs are reduced or absent from earlier diverging florideophyte lineages that secondarily simplified reproductive development, including Palmariales and Corallinophycidae. Although these patterns do not establish the ancestral functions of these genes, their evolutionary retention and diversification are consistent with a role in facilitating the evolutionary emergence of the carposporophyte as a distinct spore-producing generation, including increasingly elaborate pre- and post-fertilisation cell fusions, the proliferating gonimoblast system, and intimate physiological associations with the female gametophyte^12,15,44^.

The dependence of the carposporophyte on maternal tissues also raises broader parallels with independently evolved multicellular life cycles. In bryophytes, the diploid sporophyte remains physiologically connected to the female gametophyte through specialised transfer tissues, including the sporophytic foot, which facilitate nutrient exchange^54,55^. Similarly, in angiosperms, developing embryos depend on nutrient provisioning mediated by maternal tissues and the endosperm^56^. Although the developmental origins and tissue organisation of these systems differ fundamentally, they illustrate how sustained maternal– offspring interactions can accompany the elaboration of post-fertilisation development. Maternal provisioning may therefore represent a recurrent evolutionary strategy to the nutritional and developmental demands of increasingly complex sporophytic generations^57^.

Together, our findings support a model in which the triphasic life cycle arose though successive elaboration of an ancestral biphasic system, consistent with hypotheses based on comparative analysis of red algal life histories^45^. We propose that this transition involved three potentially interconnected processes: (1) the elaboration of ancestral post-fertilisation zygotic amplification, potentially building on the direct division of fertilised carpogonia into carpospores observed in Bangiophyceae^58,59^; (2) the evolution of maternal provisioning and coordinated but distinct developmental programs in the zygote-derived carposporophyte and surrounding female gametophytic tissues and (3) the progressive diversification of TF families and recruitment of red algal lineage-specific genes into reproductive developmental programs. Collectively, these processes provide a plausible framework for understanding how an additional multicellular generation evolved and became developmentally integrated into an ancestral biphasic life cycle. More broadly, our findings illustrate how the progressive elaboration of ancestral developmental programs, the incorporation of lineage-specific genetic innovations, and the establishment of intimate developmental interactions between generations contribute to the evolution of complex multicellular life cycles.

## Materials and Methods

### Algal culture and strains

The *Bostrychia morit*ziana strain used in this study (JW3235) was originally isolated from Beachwood, Durban, Natal, South Africa, and has been described previously^60,61^. The male and female gametophytes and tetrasporophyte were derived from the same cultures used for the *Bostrychia* reference genome assembly^19^. Algal cultures were maintained in autoclaved natural seawater (NSW) supplemented with half-strength West–McBride enrichment medium^26^. The enriched seawater was replaced weekly to maintain culture growth and health. Cultures were grown at 20 °C under a 16 h light/8 h dark photoperiod, with a light intensity of approximately 30 µmol photons m⁻² s⁻¹.

### Genomic DNA extraction and sequencing

To increase read depth for the updated genome assembly, we performed additional long-read sequencing of the male gametophyte of *Bostrychia moritziana* strain Bm3235. High-molecular-weight (HMW) genomic DNA was extracted using the same method described previously^19^. Long-read sequencing was performed on an Oxford Nanopore Technologies (ONT) PromethION instrument. Sequencing libraries were prepared using the Ligation Sequencing gDNA kit (SQK-LSK114) and run on a PromethION (FLO-PRO114M) flow cell. Basecalling was performed in duplex mode using Dorado v0.9.6 with the dna_r10.4.1_e8.2_400bps_sup@v5.0.0 configuration (https://github.com/nanoporetech/dorado). Because the *Bostrychia* genome V1 Nanopore data (BioProject PRJNA1219826) had been basecalled with an older model, we additionally re-basecalled that dataset using Dorado in simplex basecaller mode with the dna_r9.4.1_e8_sup@v3.6 configuration to ensure consistency with currently available models. Short-read sequencing data of male and female gametophytes was generated using genomic DNA extracted as described above. Sequencing libraries were prepared and sequenced commercially by GENEWIZ from Azenta Life Sciences (Leipzig, German) on an Illumina NovaSeq platform to generate 150-bp paired-end reads.

### Genome assembly

A draft genome assembly was generated using a hybrid assembly approach combining the ONT and PacBio long reads from the previous genome project (BioProject PRJNA1219826) with the newly generated ONT reads from this study, using hifiasm v0.25.0 with the -l0 and -- telo-m TTTTTAGGGG options^62^. Hi-C reads (BioProject PRJNA1219826) were incorporated for scaffolding using the --h1 and --h2 options. The draft assembly was subsequently polished using NextPolish v1.4.1, performing three rounds of polishing (rerun = 3) with short-read data of the male gametophyte aligned via BWA (sgs_options = -max_depth 100 -bwa)^63^.

To resolve the assembly at chromosome scale, Hi-C reads were used for scaffolding with a 3D-DNA de novo assembly pipeline implemented in HapHiC v1.0.7^64^. Hi-C reads were first aligned to the polished draft assembly using BWA-MEM (bwa mem -5SP), followed by duplicate marking with SAMBLASTER and filtering with SAMtools (-F 3340) to retain properly formatted alignments^65–67^. Alignments were further filtered using HapHiC’s filter_bam utility, retaining reads with a mapping quality ≥ 1 and an edit distance < 3. Scaffolding was then performed using the HapHiC pipeline command using the --quick_view mode. Subsequently, we employed Juicer to generate Hi-C contact maps and assign chromosomes^68^. Remaining assembly gaps were closed using TGS-GapCloser v1.2.1, and base-level errors were corrected using Racon v1.5.0 with both Nanopore and PacBio HiFi reads^69,70^. Chromosomes were subsequently sorted, oriented, and renamed according to the chromosome nomenclature established for *Bostrychia* genome V1.

Given the presence of bacterial communities associated with the algal thallus and consequently present in the sequencing data, we manually screened the assembly for bacterial insertions, using a combination of GC content, read-mapping coverage, and inspection of regions flanking assembly gaps to identify candidate insertions, which were removed prior to structural annotation. Telomeric repeats were identified using FindTelomeres (https://github.com/JanaSperschneider/FindTelomeres). Both the mitochondrial and plastid genomes were identified using Tiara v1.0.3^71^.

### Structural, functional and repeat annotation of *Bostrychia*

Prior to performing an updated annotation of protein-coding genes across the *Bostrychia* genome, the assembly was soft-masked for repetitive elements and transposable elements (TEs). Soft-masking was performed using RepeatMasker version open-4.0.9 with the -xsmall and -gff options, using the comprehensive, manually curated repeat library previously generated for the male genome V1 (https://www.repeatmasker.org). Tandemly repeated satellite and microsatellite elements were additionally annotated using Tandem Repeats Finder (TRF) v4.10.0, with the following parameters: match = 2, mismatch = 7, delta = 7, PM = 80, PI = 10, minscore = 50, maxperiod = 2000, and the -m -f -d -ngs flags^72^. To incorporate these tandem repeats into the soft-masked assembly, BEDTools v2.31.1 maskfasta was applied^73^.

Structural annotation of the updated male assembly was performed using two complementary approaches. First, the existing genome V1 annotation was mapped onto the new assembly V2 using Liftoff v1.6.3 with default parameters^74^. Second, a *de novo* annotation was independently generated for the V2 assembly using funannotate v1.8.17 (https://funannotate.readthedocs.io/). Briefly, RNA-seq reads from male, female, and tetrasporophyte life-cycle stages were quality-trimmed using Trim Galore v0.6.10 and used to train gene models with funannotate train, using strand-specific (--stranded RF) paired-end read alignment (https://github.com/felixkrueger/trimgalore). Gene prediction was subsequently performed with funannotate predict, incorporating protein homology evidence from a protein database derived from Eukaryota OrthoDB enriched with published red algal proteins (**Supplemental Table 6**), BUSCO-based training (eukaryota_odb10), and repeat masking (--repeats2evm)^75^. Evidence source weights were adjusted for EVidenceModeler (GlimmerHMM weight = 0, PASA weight = 10, high-quality/hint evidence weight = 4)^76^. Gene models were further refined using funannotate update, incorporating UTR annotation. Gene models from the Liftoff-based and *de novo* annotations were compared using bedtools intersect -wao to identify overlapping loci. For overlapping gene models, the V1-derived gene annotation and its gene ID were retained, for non-overlapping loci, gene models from both annotation approaches were retained independently. Gene IDs followed the standardized nomenclature established previously (Bm01g000010.t1, where “Bm” denotes *Bostrychia moritziana*, “01” the chromosome number, “g” indicates “gene,” “000010” a unique numeric gene identifier, and “.t1” the transcript number). For novel genes identified in V2 with no corresponding V1 locus, new gene IDs were assigned using the intermediate numbering steps (e.g., 000003, 000005, 000007), allowing new loci to be inserted between existing V1 gene IDs while preserving the chromosomal positional order of the original numbering scheme. To simplify downstream analyses, only the longest transcript isoform per gene was retained. The gene set was further filtered to exclude TE-related genes, defined as genes with ≥95% of their coding sequence overlapping soft-masked repetitive regions. Remaining genes were then classified as either protein-coding or putative based on the presence of at least one of the following supporting lines of evidence: transcript evidence from RNA-seq data, a functional annotation assignment, or sequence homology identified via BLAST against the red-algal-enriched Eukaryota OrthoDB protein database.

Long non-coding RNA (lncRNA) annotation was performed by integrating transcriptome assembly with coding potential classification. RNA-seq reads from multiple tissues and life-cycle stages were aligned to the V2 genome assembly using HISAT2 v2.2.1 with default parameters^77^. Transcripts were assembled for each sample using StringTie v2.1.7^78^ and individual sample assemblies were subsequently merged into a single non-redundant transcript set using stringtie --merge, followed by comparison against the reference annotation using GffCompare v0.12.10 to assign class codes (https://gffutils.readthedocs.io/). Candidate lncRNAs were then identified using the FEELnc pipeline v0.2.1 using default parameters^79^.

Functional and protein domain annotation was conducted using InterProScan v5.64-96.0 and EggNOG-mapper v2.1.8^80–82^. A curated annotation of transcription-associated proteins (TAPs) was carried out using TAPScan v4^83^. Gene prediction, functional annotation, as well as rRNA and tRNA annotation of the plastid genome was performed using GeSeq^84^. The same annotations, including gene prediction, functional annotation, and rRNA and tRNA annotation, were performed for the mitochondrial genome using MFannot^85^.

### Structural gene annotation of published red algal genomes

For comparative genomic analyses, gene annotations were additionally generated for eleven published red algal genome assemblies lacking existing annotations: *Amphiroa fragilissima*, *Gulsonia nodulosa*, *Lithothamnion corallioides*, *Osmundea hybrida*, *Phymatolithon calcareum*, *Eucheuma denticulatum*, *Kappaphycus striatus*, *Palmaria palmata*, *Pyropia vietnamensis*, *Galdieria sulphuraria*, and *Galdieria partita*. For each assembly, repetitive elements were identified *de novo* using RepeatModeler v2.0.7 and soft-masked using RepeatMasker together with TRF, following the same repeat-masking approach described for *Bostrychia*^86^. Publicly available RNA-seq reads (where available) were quality-trimmed with Trim Galore and aligned to each masked genome using HISAT2. Gene models were then predicted using BRAKER3, incorporating RNA-seq evidence and a red-algal-enriched Eukaryota OrthoDB protein database, followed by removal of TE-related gene models as described for *Bostrychia*^87–92^.

### RNA-seq analysis

RNA-Seq libraries were generated from low input total RNA isolated from dissected *Bostrychia* thallus, isolated spores and spermata using Smart-seq3 with independent biological replicates^93^. Total RNA was isolated using an RNeasy PowerPlant Kit (QIAGEN) and followed by ethanol precipation to concetrate RNA prior to library preapration. The libraries were sequenced on an Illumina NovaSeq 2000 using 150-bp paired-end reads. RNA-seq reads were processed with nf-core/rnaseq v3.12.0 using Nextflow v23.10.0^94^. Adapter and quality trimming was performed with Trim Galore v0.6.7 (https://github.com/FelixKrueger/TrimGalore). Reads were aligned to the chromosome-scale *Bostrychia* genome V2 (this study) using STAR v2.7.9a/v2.7.10a, and transcript abundance quantified with Salmon v1.10.1^95,96^. Alignment files were processed using SAMtools v1.16.1/v1.17, while StringTie v2.2.1 was used by the workflow for transcript-level processing^78,97^. The downstream analysis used merged gene-level count and transcripts-per-million (TPM) matrices produced by the STAR-Salmon workflow. Gene models were restricted to biotype ‘protein_coding’, yielding a total 18,714 protein-coding genes. For count-based analyses, genes were filtered with *filterByExpr* in edgeR v3.42.4, using developmental stage as the grouping factor^98^. This retained 13,285 detectably expressed protein-coding genes. TPM-based summaries retained the full protein-coding universe to distinguish absence of detectable expression from exclusion during count filtering.

Sample relationships were assessed by principal-component analysis of regularised-logarithm-transformed counts generated with DESeq2 v1.40.2^99^. Gene-body coverage profiles and 5’ to 3’ coverage ratios were obtained from Qualimap summaries^100^. Smart-seq3 profiles from this study were compared with previously published *Bostrychia* thallus libraries^19^. The latter were not included in any differential expression, marker, co-expression or enrichment analysis. Differential expression analysis was performed using DESeq2 v1.40.2 using the count-filtered protein-coding gene matrix^99^. Counts were rounded to integers, genes with zero counts in all included samples were removed, and a single model with developmental stage as the explanatory variable (‘design = ∼ stage’) was fitted across the 42 libraries. Pairwise contrasts were then extracted from this fitted model for stage comparisons. Unless stated, differentially expressed genes were defined by an absolute log2 fold change of at least 1 and a Benjamini-Hochberg false-discovery rate (FDR) ≤ 0.05.

To test whether broad developmental expression patterns depended on highly variable genes during library normalization, an alternative set of DESeq2 size factors was estimated from low-variance control genes. Genes detected above the TPM threshold in every developmental stage were ranked by variance, and the 724 lowest-variance genes among 8,828 ubiquitously detected genes were supplied to DESeq2 as *controlGenes*. Differential expression was then recalculated using these alternative size factors. A separate conservative analysis restricted differential-expression testing itself to the same 724-gene subset. Robustness was evaluated for both the fraction of genes classified as stage dependent and the dynamic range of genome-wide expression. Dynamic range was calculated for each gene as the difference between its maximum and minimum log_2_-transformed stage-mean normalized expression.

### Allele-specific expression analysis

Allele-specific expression was assessed from fixed sequence differences between the male and female parental genomes. Candidate diagnostic variants were obtained from a joint variant call set processed with nf-core/sarek v3.5.1 using short-read data of the male and female gametophyte^94^. Sites were retained when both parents were homozygous for discordant alleles, parental depth was at least 12, genotype quality was at least 30, the minor-allele fraction within each parent was at most 0.05, and no indel occurred within 5 bp. Variants on the U and V sex chromosomes were excluded. An N-masked reference was generated at diagnostic sites and RNA-seq reads were realigned with STAR v2.7.11b^95^. Alignments were split by allele with SNPsplit v0.6.0 and assigned to genes with HTSeq v0.11.2^101,102^.

### Stage-specific and stage-marker genes

Stage specificity was first summarized from mean TPM values calculated across biological replicates. For each gene, tissue specificity was quantified using *tau*, calculated as the sum of ‘1 - x_i/max(x)ߣ across stages divided by the number of stages minus one^103^. Strict stage markers were required to have *tau* ≥ 0.7, a maximum stage-mean abundance of TPM > 5, detectable expression of at least 1 TPM in no more than four stages, and support from a relevant differential-expression contrast involving the candidate stage. The differential-expression requirement was an oriented log_2_ fold change ≥1 in favour of the candidate stage and FDR ≤ 0.05. Conversely, stage-unique genes were defined as genes with TPM > 1 in only one of the 13 stages.

### Generation-biased and generation-repressed genes

Generation identity was assessed by grouping samples according to their generation of origin, excluding reproductive cells and tissues containing contributions from more than one generation. The gametophyte group comprised female vegetative tissue, procarp-bearing female tissue, male vegetative tissue and spermatangial tissue but excluded spermatia. The carposporophyte group included only isolated carposporophytes, excluding pericarp and whole cystocarps. The tetrasporophyte group comprised vegetative tissue and stichidia. Counts were modelled with DESeq2 after assigning samples to the three generation groups. For each group, biased genes were defined as the intersection of genes significantly upregulated relative to each of the other two groups, requiring a log_2_ fold change ≥ 1 and FDR ≤ 0.05 in both comparisons. Repressed genes were similarly defined as the intersection of genes significantly downregulated relative to both other groups, requiring a log_2_ fold change ≤ -1 and FDR ≤ 0.05.

### Sex-biased expression and UV sex chromosome analyses

Sex-biased expression was examined within the haploid gametophytes using a pooled female-versus-male contrast. The female group comprised female vegetative and procarp-bearing samples, whereas the male group comprised male vegetative and spermatangial samples. Spermatia were excluded to avoid confounding sex with a highly differentiated reproductive-cell transcriptome. Female-biased genes were defined by a log_2_ fold change ≥ 1 and FDR ≤ 0.05, and male-biased genes by log_2_ fold change ≤ -1 and FDR ≤ 0.05. Genes were classified as autosomal, U-linked, V-linked or pseudoautosomal using the *Bostrychia* v2 genome annotation (this study). Mitochondrial and plastid chromosomes were excluded. Enrichment of sex-biased genes in each genomic compartment was tested using two-sided Fisher’s exact tests followed by Benjamini-Hochberg correction. The UV expression heatmap included all protein-coding genes on these chromosomes, grouped by chromosome and pseudoautosomal or sex-determining region, with genes clustered by expression within each group. Cells corresponding to an SDR absent from the sampled haploid sex were masked.

### Weighted gene co-expression network analysis

Weighted gene co-expression network analysis was performed with WGCNA v1.72.5 ^104^. The analysis used the 13,285 protein-coding genes retained by *filterByExpr* and transformed as ‘log_2_(count + 1)’ across all 42 included libraries. Genes and samples failing the WGCNA *goodSamplesGenes* check were excluded. A signed network and signed topological overlap matrix were constructed with a soft-thresholding power of 12. Modules were inferred using *blockwiseModules* with the following paramters: *deepSplit = 2*, *minModuleSize = 30*, *mergeCutHeight = 0.2*, *pamRespectsDendro = FALSE, reassignThreshold = 0, maxBlockSize = 5000*. This assigned 13,078 of the 13,285 filtered genes to 23 co-expression modules, with the remaining 207 genes forming an unassigned class. Soft-thresholding powers of 1–20 were examined using signed-network scale-free topology diagnostics. The final network used a power of 12, for which the archived diagnostic gave an R-squared of approximately 0.70. Module-detection parameters were subsequently evaluated over *deepSplit* values of 2-4, minimum module sizes of 20 or 30 and merge heights of 0.10, 0.15 or 0.20. The final parameter set was chosen to retain biologically relevant and meaningful stage profiles. Module eigengenes were calculated as the first principal component of each module’s expression matrix. Stage profiles were obtained by averaging eigengene values across replicates and standardizing these means across stages within each module. Modules were hierarchically clustered from the standardized stage profiles using Euclidean distance and Ward’s D2 linkage. The 24 WGCNA colour classes were renamed M01-M24 according to their order in this dendrogram. M21 corresponds to the grey unassigned class and thus does not represent a coherent co-expression module. Gene-module membership was quantified as signed eigengene connectivity (kME), and hub genes were defined as the top 10% of genes ranked by absolute kME within each assigned module.

### Functional enrichment analysis

Of the 18,714 protein-coding genes, 10,221 had at least one Gene Ontology (GO), KEGG Orthology, Pfam and COG assignment. Enrichment was tested using one-sided Fisher’s exact tests, with the relevant full protein-coding universe as the background. Terms represented by fewer than five background genes were generally excluded, with a minimum of three genes used for the smaller stage-unique sets. P values were corrected by the Benjamini-Hochberg procedure. For module enrichment, modules containing at least ten genes were tested. The high-confidence module terms required FDR ≤ 0.05, at least three overlapping genes, an odds ratio ≥ 2 and a background term size of 10-500 genes. Broad ontology and generic parent terms were removed from the summarized output. Generation-biased and generation-repressed sets were tested for GO and KEGG enrichment. Benjamini-Hochberg correction was applied jointly across the GO and KEGG tests for the three generation-biased sets, and separately across the corresponding tests for the three generation-repressed sets. These global adjusted P values are reported as FDR in Supplemental Table 5. Additional within-set adjusted P values were calculated separately for each generation-identity set and retained in the analysis outputs. Sex-biased and marker-gene enrichments were assessed using the same Fisher-test framework with the annotation classes available for each analysis. TAP enrichment was tested using one-sided Fisher’s exact tests for co-expression modules and two-sided Fisher’s exact tests for generation-identity and sex-biased gene sets, followed by Benjamini-Hochberg correction. For the module lollipop plots, all module genes with a TAP-family annotation were ranked by absolute kME and the 15 highest-ranking TAPs in each module plotted.

### Phylostratigraphy and transcriptome age index

Gene ages were inferred using genERA^105^, which assigned phylostratigraphic age to 18,318 protein-coding genes. Transcriptome age index (TAI) was calculated using myTAI v2.2.0.9006 as the expression-weighted mean phylostratum rank at each developmental stage, using replicate-averaged TPM values^34^. Sensitivity analyses used square-root, log₂ and rank transformations of expression values, together with removal of genes above selected quantiles of mean expression. Transformations producing negative or non-finite expression weights were excluded from interpretation as TAI estimates. Deviation from a developmentally flat age profile was assessed using the myTAI *flat-line* test. Enrichment or depletion of individual phylostrata within gene sets was evaluated with one-sided Fisher’s exact tests followed by Benjamini-Hochberg correction. Module-level shifts in the complete phylostratigraphic composition were additionally assessed using chi-squared tests, with significance verified against 20,000 simulated tables. Organelle genes were excluded from chromosome-based phylostratigraphic summaries.

### Chromosomal clustering of co-expressed and identity genes

Spatial clustering was tested for WGCNA modules, the six generation-biased or generation-repressed sets, and female- and male-biased genes. Each gene was represented by the midpoint of its annotated genomic interval. For every gene in a tested set, the distance to its nearest same-set gene on the same chromosome was calculated, and the observed set statistic was the median nearest-neighbour distance. The null distribution was obtained from 1,000 chromosome-stratified permutations that preserved both gene set size and the number of selected genes on each chromosome. Mitochondrial and plastid chromosomes were excluded. Analyses used a fixed random seed of 42. Empirical one-sided P values were calculated as (1 + number of permuted medians ≤ observed median)/(1 + number of permutations) and corrected across gene sets using the Benjamini-Hochberg method. A set was classified as significantly clustered when FDR ≤ 0.05 and its standardized distance statistic was negative. The expected distance was defined as the median of the median nearest-neighbour distances obtained across permutations. Generation-identity plots show log₂(observed/expected distance), with negative values indicating shorter distances than expected. Module clustering-strength plots show the reciprocal ratio, log₂(expected/observed distance), with positive values indicating stronger clustering. To define interpretable physical clusters, adjacent set members on the same chromosome were joined when their gene midpoints were separated by no more than 50 kb. Blocks containing at least two genes were retained and assigned stable cluster identifiers.

### Comparisons of module connectivity

To explore whether fixed co-expression gene sets showed similar internal connectivity across developmental stages, connectivity was recalculated separately within each stage. The 23 assigned WGCNA modules and the grey unassigned class were retained as fixed gene sets. Gene–gene Pearson correlations were calculated from log₂(count + 1) values across the libraries within each stage, using six libraries for the carposporophyte and three for each other stage. Undefined correlations were set to zero. Signed adjacency was calculated as ((1 + r)/2)¹², and intramodular connectivity was obtained by summing adjacencies to the other genes in the set. Similarity to the isolated carposporophyte was quantified as the Spearman correlation between gene-connectivity ranks.

### Analysis of module and TF evolution

To analyze module and TF evolution, protein sequences of 38 species of red algae and the green lineage were grouped into orthogroups using OrthoFinder v2.5.5^106^. For this, publicly available proteomes of red algae and the green lineage were combined with proteomes from newly annotated red algae genomes generated in this study. The rooted species tree inferred by OrthoFinder was used as the phylogenetic backbone for all downstream analyses, rooted on *Guillardia theta*. Genes assigned to each of the 24 co-expression modules were mapped to their corresponding orthogroups using the gene-to-orthogroup assignments in Orthogroups.tsv. Where multiple genes within a module were assigned to the same orthogroup, the orthogroup was counted once per module, such that each module was represented by its underlying set of distinct orthogroups. To trace the evolutionary origin of each module’s gene complement, orthogroup presence/absence across all species was reconstructed at every internal node of the rooted species tree using Dollo parsimony, implemented in Count v9.1106, which assumes a single origin per orthogroup and allows independent losses along the tree^107^. For each module, per-node module completeness was calculated as the proportion of the module’s constituent orthogroups reconstructed as present at that node. For the TFs most strongly associated within each module, the same per-orthogroup Dollo reconstruction was used to identify each TF’s node of origin.

### Identification of paralogous transcription factors

To assess whether TFs within a given co-expression module represent duplicated paralogs or evolutionarily unrelated members of the same TF family, we combined orthogroup assignment with duplication inference from OrthoFinder. The OrthoFinder run generated hierarchical orthogroups (HOGs) at each node of the rooted species tree, alongside a genome-wide set of inferred gene duplication events (Duplications.tsv). For each TF gene, HOG membership was extracted at the node immediately preceding the divergence of *Bostrychia* from its closest relatives in the species tree. TF genes co-occurring within a module and TF family were classified as duplicated (paralogs) when they shared this HOG and were linked by at least one inferred duplication event, all other TF genes were considered evolutionarily unrelated members of the same TF family. For each duplicated TF family, the phylogenetic origin of the underlying orthogroup and the position of the corresponding duplication event were mapped onto the species tree.

## Acknowledgments

We thank Javier Lazaro for scientific illustration of the *Bostrychia* life cycle. R.P. is a member of the International Max Planck Research School “From Molecules to Organisms”. This work was supported by the Max-Planck-Gesellschaft.

## Author contributions

Conceptualization, M.B.; methodology, R.P., P.S., V.B and M.B.; investigation, R.P., P.S. and M.B.; visualization, R.P. and M.B.; resources, J.A.W. and S.M.C.; data curation, M.B..; writing – original draft, M.B.; writing – review and editing, R.P., S.M.C. and M.B.; supervision, M.B.; project administration, M.B.

## Data and code availability

The raw Oxford Nanopore Technologies (ONT) long-reads and Illumina DNA and RNA sequencing (RNA-seq) short-reads have been uploaded to BioProject PRJNA1555016. A repository containing phylogenetic trees, the re-annotated red algal genomes used in this study, as well as the *Bostrychia moritziana* v2.0 genome assembly and its associated annotation can be accessed at https://doi.org/10.57892/100-380. All original code is publicly available at https://github.com/Borg-Lab/triphasic-lifecycle-analysis. Any additional information required to reanalyze the data reported in this paper is available from the corresponding author upon request.

## Supplemental Tables

Supplemental Table 1 – Structural annotation of the *Bostrychia* v2 genome

Supplemental Table 2 – RNA-seq mapping statistics

Supplemental Table 3 – Gene expression values and gene annotation

Supplemental Table 4 – Statistical analysis of chromosomal clustering

Supplemental Table 5 – Statistical analysis of functional enrichment

Supplemental Table 6 – Red algal and outgroup species used in this study

Supplemental Table 7 – Transcription-associated protein (TAP) evolution across co-expression modules

---

**Extended Data Figure 1.**
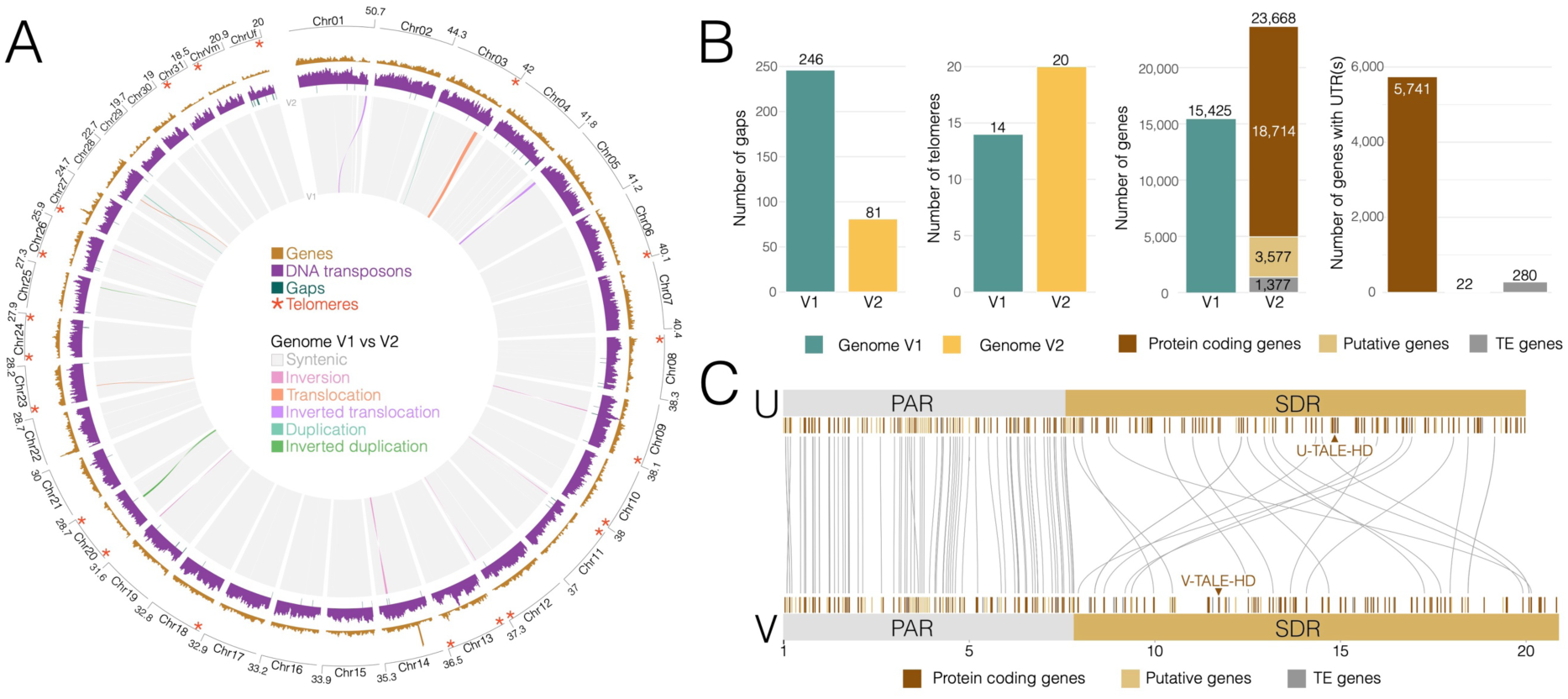
Improved genome assembly and annotation of *Bostrychia moritziana*. (A) Circos plot comparing the previous (V1) and updated (V2) genome assemblies. Chromosomes are arranged around the outer ring, with chromosome lengths (Mb), gene density (orange), DNA transposon density (purple), assembly gaps (green) and identified telomeric repeats (red asterisks) indicated. Ribbons connecting genomic regions represent syntenic blocks and structural differences between assemblies, including inversions, translocations and duplications, as indicated by the colour key. (B) Comparison of assembly and annotation statistics between V1 and V2, showing the number of assembly gaps, chromosome ends with identified telomeric repeats, annotated genes and genes with annotated untranslated. (C) Comparison of the U and V sex chromosomes in the updated assembly, showing the pseudoautosomal regions (PARs) and sex-determining regions (SDRs). Vertical bars indicate annotated genes, coloured according to their classification, and grey connecting lines indicate homologous genes between the U and V chromosomes. The positions of the U- and V-linked TALE homeodomain transcription factors (U-TALE-HD and V-TALE-HD) are indicated.

**Extended Data Figure 2.**
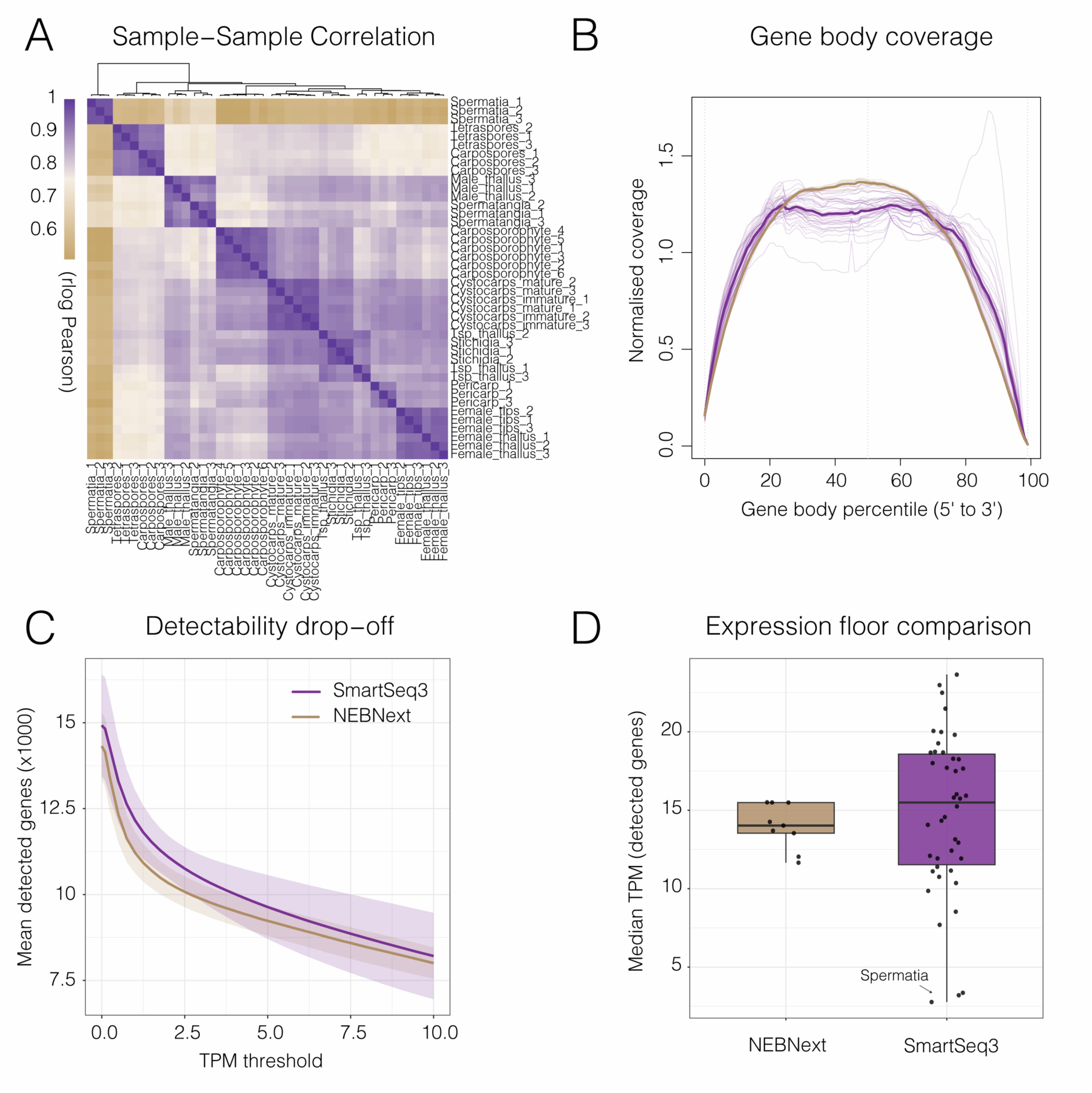
RNA-seq quality assessment and transcript detectability of Smart-seq3 libraries. (A) Heatmap of pairwise Pearson correlations (log-transformed TPM values) across RNA-seq samples. (B) Normalised gene body coverage across transcripts, from the 5′ to the 3′ end. Individual lines represent samples, with thicker lines indicating average coverage profiles for Smart-seq3 (purple; this study) and NEBNext Ultra II libraries (brown; ^19^). (C) Transcript detectability across increasing expression thresholds (TPM), showing the mean number of detected genes for each library type. Shaded regions indicate ±1 standard deviation. (D) Distribution of median TPM values among detected genes (TPM > 0) for individual samples, grouped by library preparation method. Points represent individual samples, with spermatia highlighted as having particularly low median expression values.

**Extended Data Figure 3.**
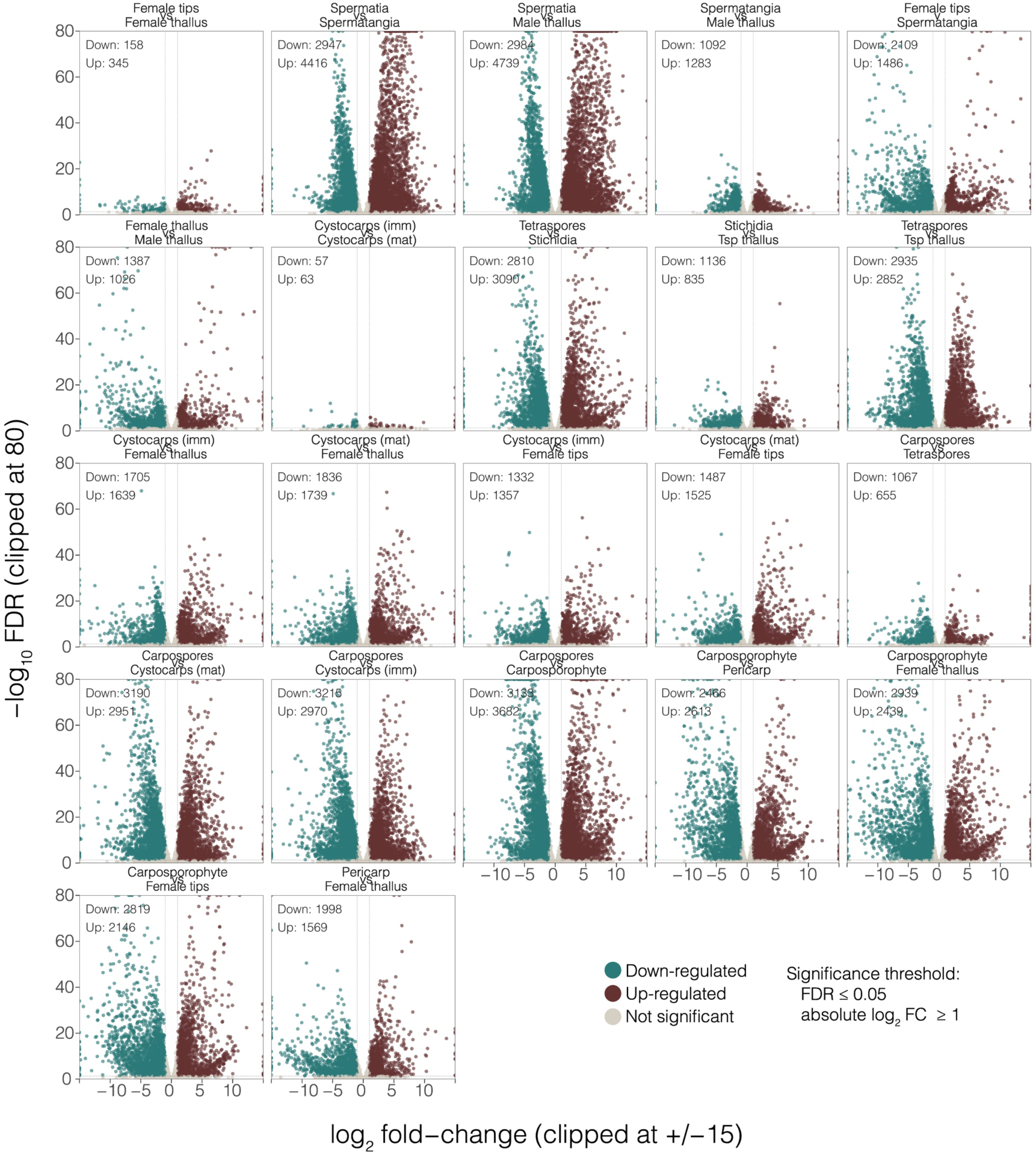
Differential gene expression across the *Bostrychia* life cycle. Volcano plots show pairwise differential gene expression between vegetative tissues, reproductive structures and developmental stages, including gametophytes, tetrasporophytes, carposporophytes, pericarps, cystocarps and reproductive spores. Significantly differentially expressed genes (FDR ≤ 0.05 and |log₂ fold change| ≥ 1) are coloured in teal (downregulated) or maroon (upregulated), relative to the first tissue or stage named in each comparison. Non-significant genes are shown in grey. Imm, immature; mat, mature; Tsp, tetrasporophyte.

**Extended Data Figure 4.**
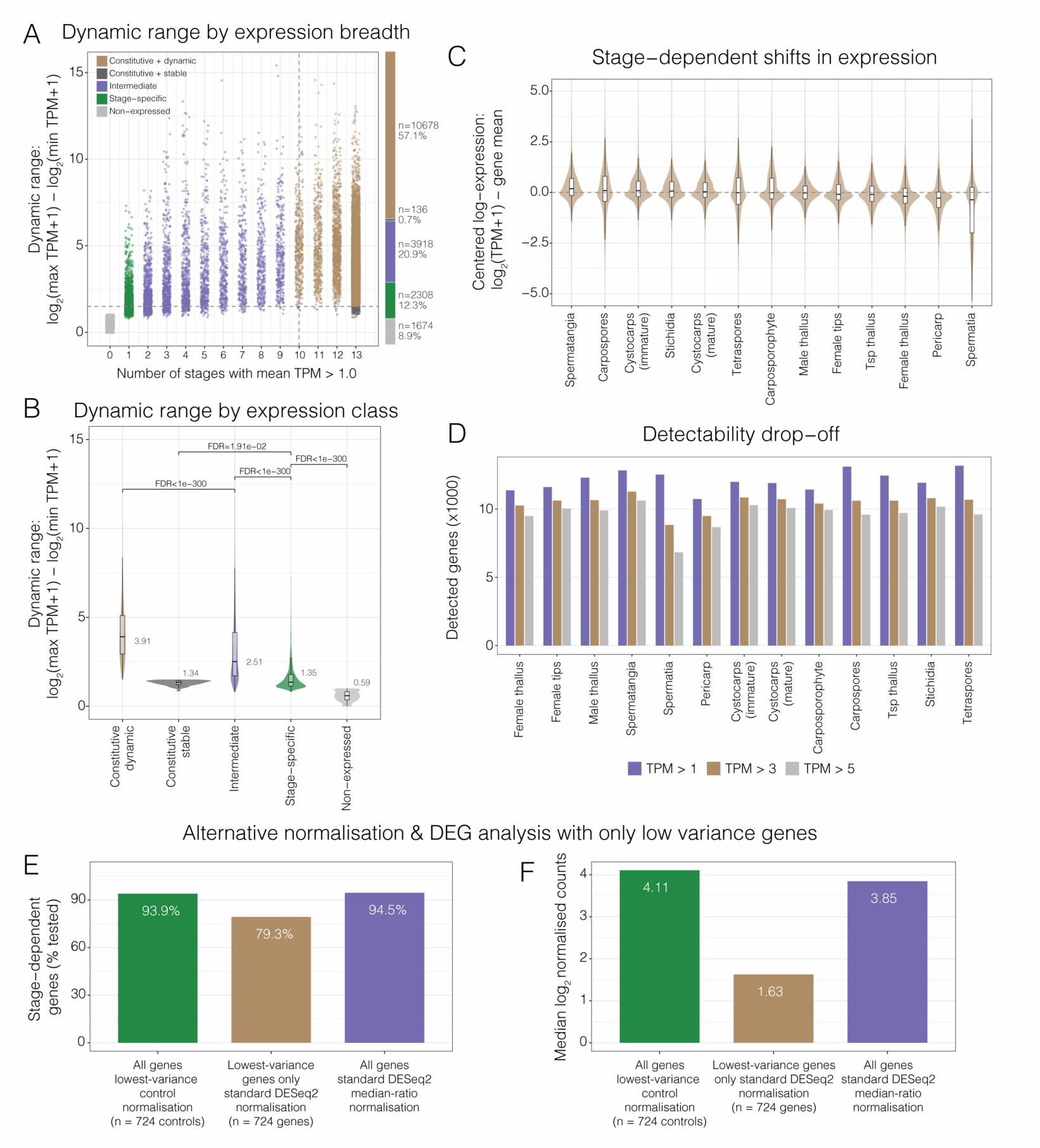
Transcriptional variation across the *Bostrychia* transcriptomic datasets. (A) Relationship between the breadth of gene expression (defined as the number of developmental stages with mean expression >1 TPM), and dynamic range (calculated as log₂(maximum TPM + 1) − log₂(minimum TPM + 1)). Genes are classified as constitutive and dynamic, constitutive and stable, intermediate, stage-specific or non-expressed, with the number and proportion of genes in each class indicated alongside the bar chart. (B) Distribution of the dynamic range in expression across the five expression classes, with median values and false discovery rate (FDR)-adjusted significance levels indicated. (C) Distribution of gene-centred log₂(TPM + 1) expression values across developmental stages, illustrating stage-dependent deviations from each gene’s mean expression level. (D) Number of genes detected in each developmental stage or tissue at expression thresholds of TPM >1, >3 and >5. (E, F) Sensitivity analysis comparing alternative DESeq2 normalisation strategies using all genes or the 724 genes with the lowest expression variance as controls. (E) Bar chart showing the proportion of genes classified as stage-dependent under each normalisation strategy. (F) Median log₂-normalised gene counts under the corresponding strategies. These analyses assess the robustness of stage-dependent expression patterns to alternative normalisation approaches.

**Extended Data Figure 5.**
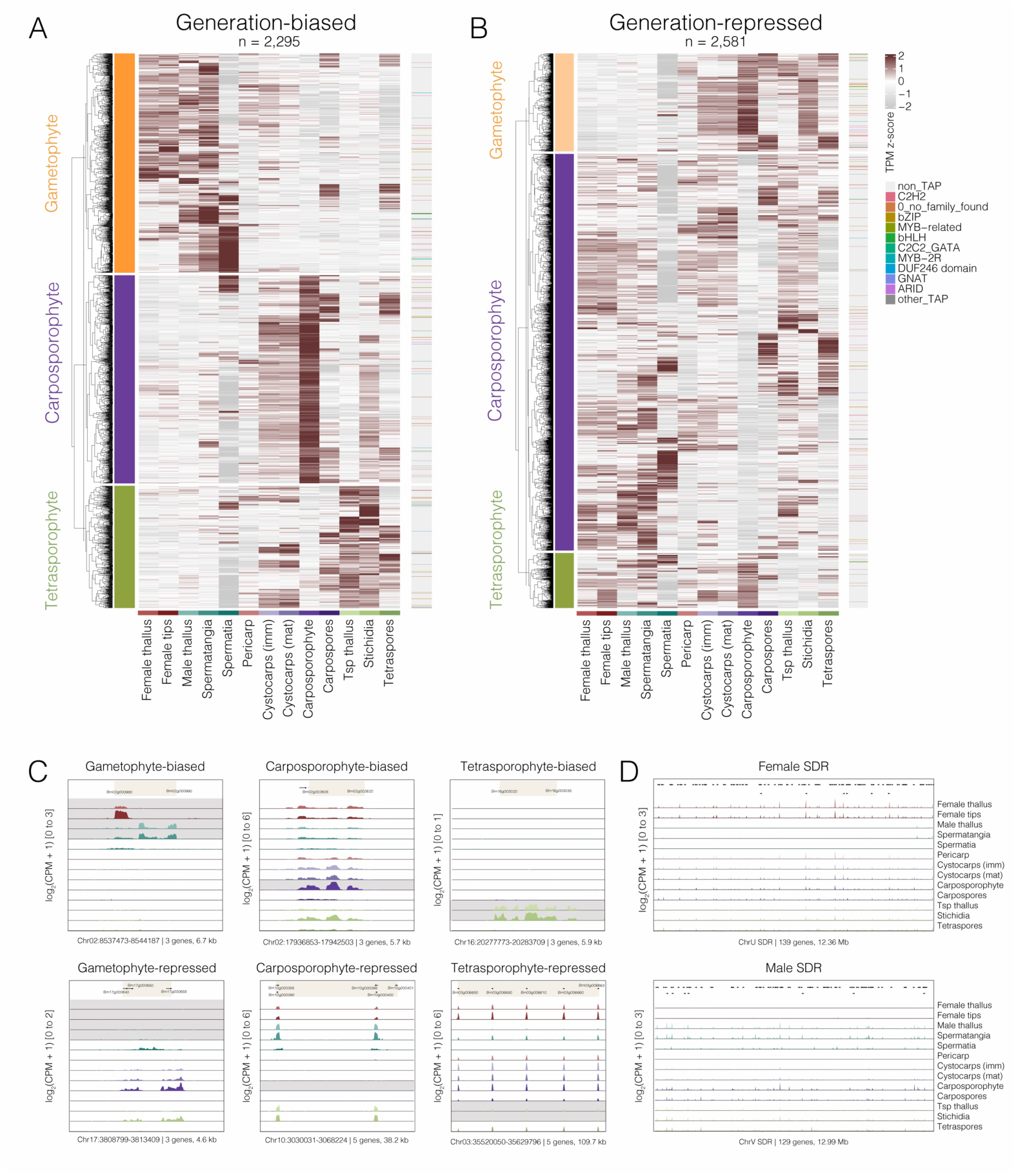
Generation-biased and generation-repressed gene expression in *Bostrychia*. (A,B) Heatmaps showing the expression of genes classified as generation-biased (n = 2,295; A) or generation-repressed (n = 2,581; B) across reproductive stages and tissues. Genes are hierarchically clustered and grouped according to the generation in which they show elevated or reduced expression, respectively. Colours represent row-scaled TPM expression values (z-scores). The coloured annotation strip on the right indicates transcription-associated protein (TAP) families. (C) Representative genome browser tracks showing clustered genes with generation-biased (top) or generation-repressed (bottom) expression in gametophytes, carposporophytes and tetrasporophytes. Tracks show log₂(CPM + 1) expression across reproductive stages and tissues, illustrating the genomic proximity of genes with similar generation-associated expression patterns. Genomic coordinates, the number of genes in each cluster, and region lengths are indicated beneath each panel. (D) Gene expression across the female (U) and male (V) sex-determining regions (SDRs). Tracks show log₂(CPM + 1) expression across reproductive stages and tissues, with annotated gene positions indicated above each panel. Genomic coordinates and the number of genes within each SDR are shown below.

**Extended Data Figure 6.**
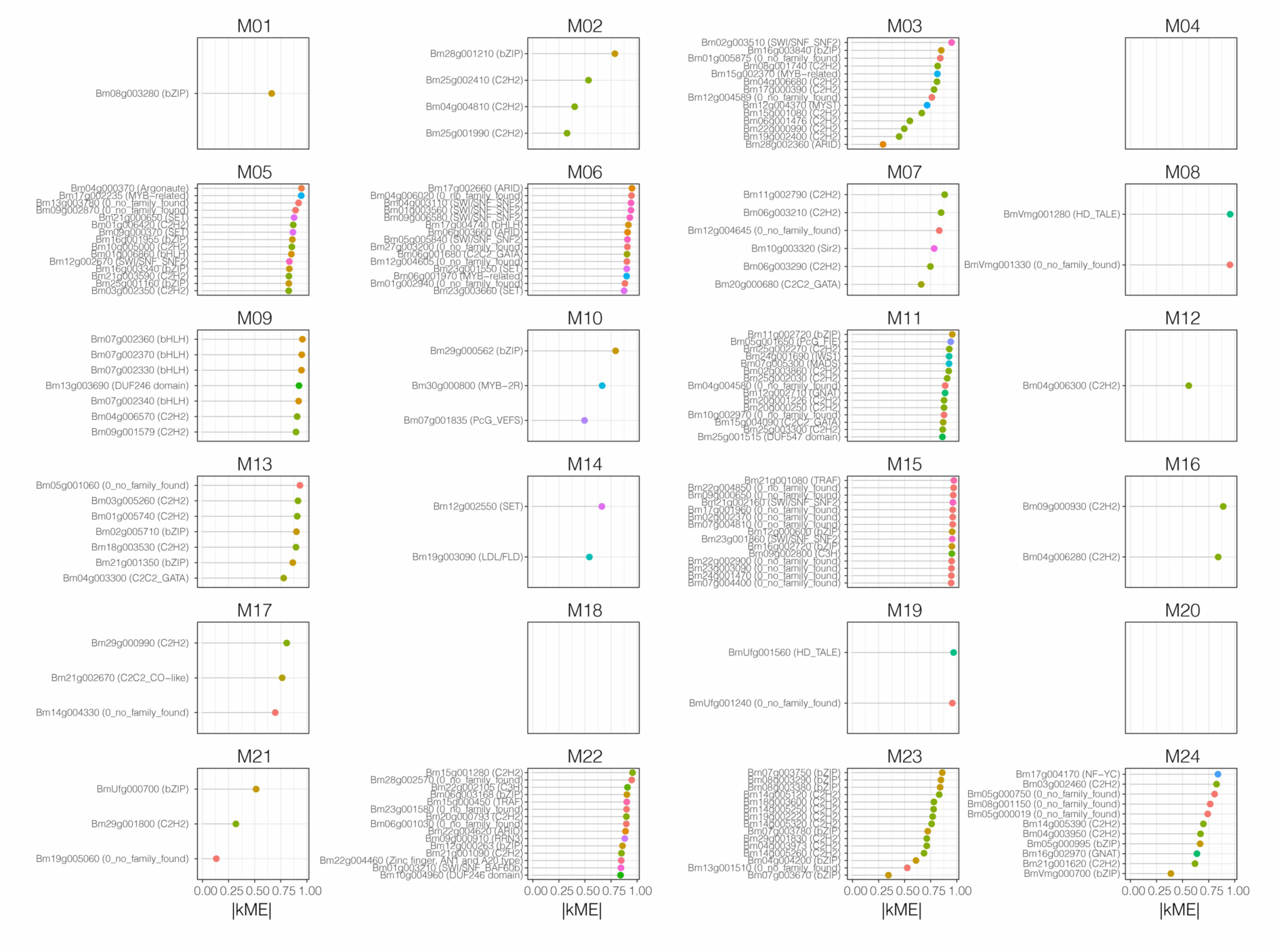
Top transcription-associated proteins (TAPs) associated with each co-expression module. For each module, TAPs are ranked by their absolute module eigengene correlation, |kME|, which measures how closely the expression profile of each TAP follows the module eigengene. Colours denote TAP family, while facets are ordered by module.

**Extended Data Fig. 7.**
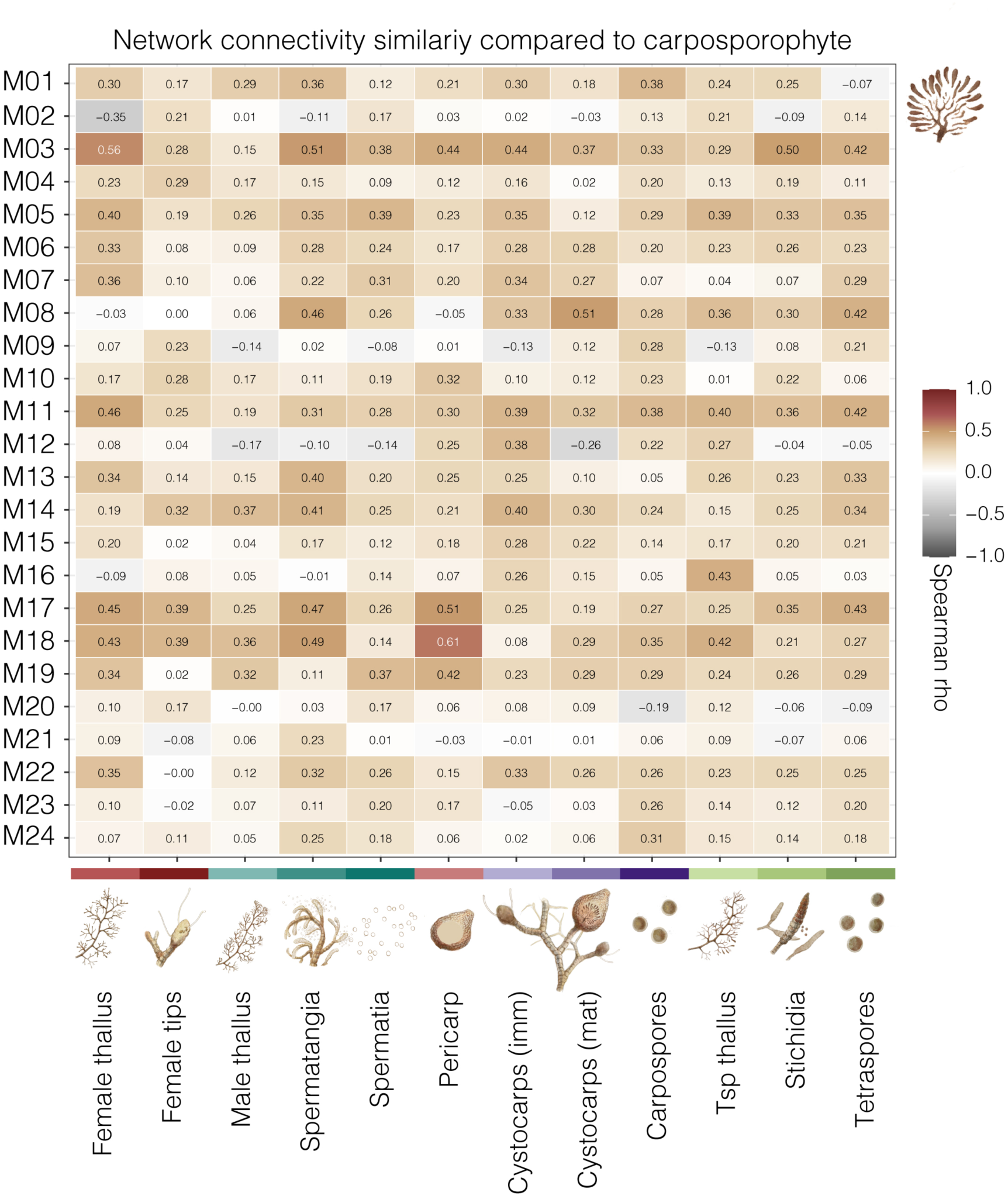
Network connectivity comparison of co-expression modules relative to the carposporophyte. Connectivity was calculated separately within each stage using the fixed WGCNA modules. Values show Spearman correlations between gene-connectivity ranks in the isolated carposporophyte and each comparison stage, with the carposporophyte self-comparison omitted.

**Extended Data Fig. 8.**
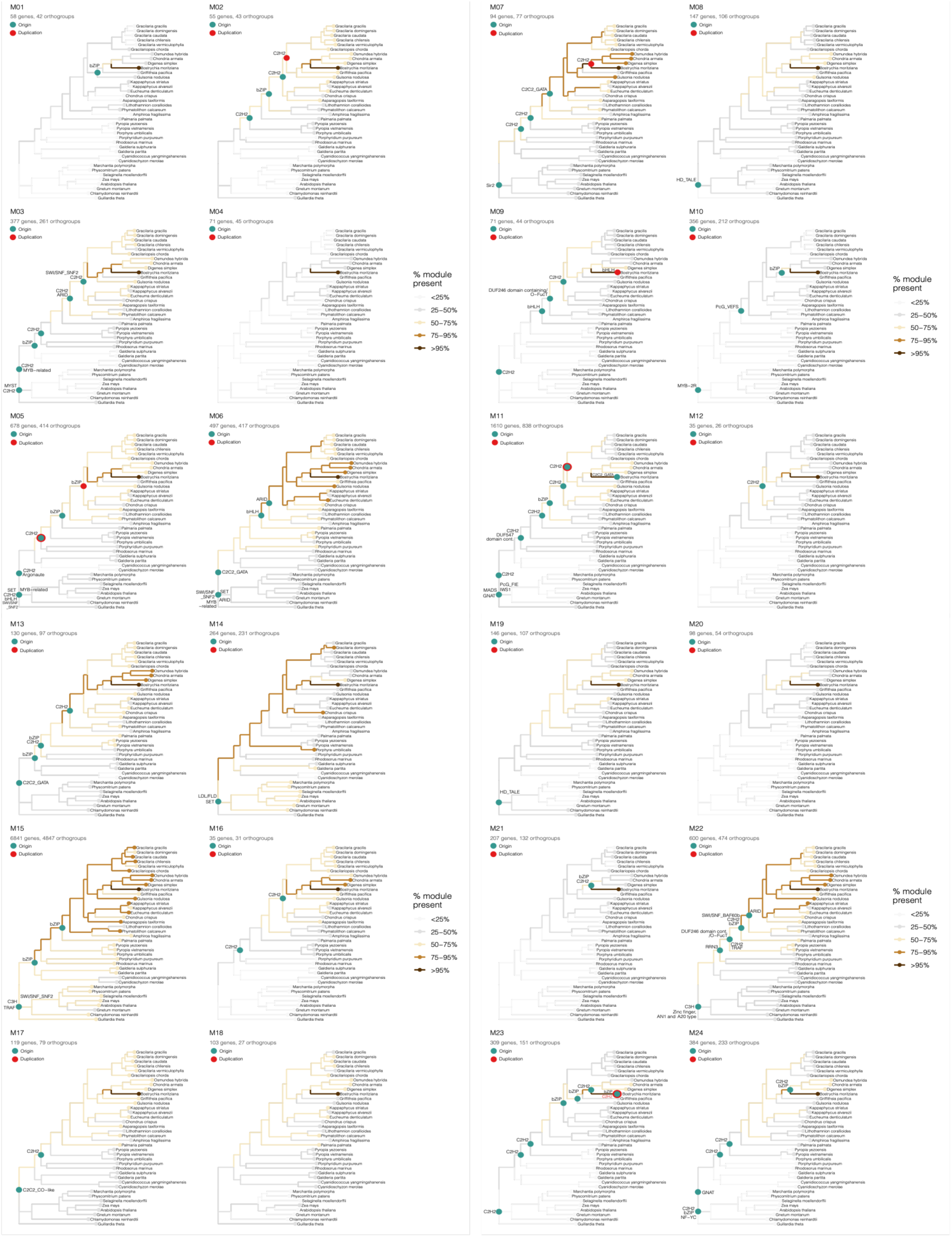
Phylogenetic distribution and inferred evolutionary histories of orthogroups associated with the co-expression modules. Branch colours indicate the proportion of module-associated orthogroups present in each lineage, while coloured nodes indicate inferred origins (teal) and duplications (red) of selected TF families. Imm, immature; mat, mature; Tsp, tetrasporophyte; TPM, transcripts per million.

